# Gestational Chronic Intermittent Hypoxia Triggers Maternal Inflammation and Disrupts Placental Stress Responses

**DOI:** 10.1101/2025.05.29.656944

**Authors:** Jennifer J. Gardner, Reneé de Nazaré Oliveira da Silva, Jessica L. Bradshaw, Steve Mabry, E. Nicole Wilson, Nataliia Hula, Selina M. Tucker, Isabelle K. Gorham, Desirae Escalera, Leslie Lopez, Nicole R. Phillips, Rebecca L. Cunningham, Styliani Goulopoulou

**Author notes:** **Correspondence:** Styliani Goulopoulou, Ph.D., Lawrence D. Longo, MD Center for Perinatal Biology, Department of Basic Sciences, Loma Linda University School of Medicine, CA, USA. Equal contribution (Jennifer J. Gardner, Reneé de Nazaré Oliveira da Silva).

## Abstract

Gestational hypoxia is associated with placental cellular responses, including oxidative stress and inflammation. Circulating cell-free mitochondrial DNA (ccf-mtDNA) is a marker of cell stress, that can be transported within extracellular vehicles (EVs), eliciting proinflammatory responses. We hypothesized that systemic exposure to chronic intermittent hypoxia (CIH) during late pregnancy would increase maternal inflammation, alter circulating EV characteristics, and disrupt placental stress responses. Pregnant rats were exposed to CIH (n=8) or normoxia (n=9) during gestational days 15-20 (term 22-23 days). On GD20, ccf-mtDNA and EV-associated mtDNA (EV-mtDNA) were quantified with qRT-qPCR, while maternal circulating cytokines were quantified using a MILLIPLEX® cytokine array. Systemic oxidative stress was measured by plasma advanced oxidation protein products (AOPP). Placental stress responses were evaluated by examining the balance between proinflammatory and antioxidant gene expression and the activation of proteins involved in apoptotic and autophagic processes. CIH exposure increased placental weights (p=0.015) and reduced placental efficiency (p=0.0006) without affecting fetal biometrics (p>0.05). Absolute ccf-mtDNA and EV-mtDNA content were unchanged (p>0.05), but EV concentrations were reduced (p=0.011) in response to CIH, suggesting an increase in EV-mtDNA per EV. Maternal interleukin-18 (IL-18) concentrations increased in the CIH group (p=0.047). Placental mRNA expression of *catalase* (p=0.048) and *sod2* (p=0.038) were upregulated, while autophagy-related proteins Beclin-1 (p=0.006) and p62 (p=0.023) were also increased in response to CIH, with no changes in LC3A/B expression (p>0.05). Gestational CIH disrupts maternal EV and inflammatory profiles, reduces placental efficiency, and modulates placental antioxidant and autophagic mechanisms, without impairing fetal growth in rats.

## Introduction

Systemic and placental hypoxia during pregnancy is associated with maternal and fetal complications, such as intrauterine growth restriction (1) and preeclampsia (2), which affect approximately 2-8% (3) and 5-10% (1) of pregnancies, respectively. These complications are also common in conditions that expose the mother and fetus to intermittent or sustained hypoxemia, including maternal smoking (4, 5), asthma (6), anemia, COVID-19 infection, high-altitude exposure, and sleep-disordered breathing (1, 7), the latter affecting an estimated 4-27% of pregnant women (8).

Physiological adaptations to hypoxia during pregnancy involve tightly coordinated maternal and fetal adaptations, with the placenta playing a central role in mediating oxygen and nutrient exchange. In early pregnancy, hypoxia is a physiological state that supports embryogenesis and organogenesis by protecting effect against oxidative stress (9–11). As gestation progresses and uteroplacental blood flow is established (∼10-12 weeks of gestation), tissue oxygenation increases, although it remains lower than adult levels (10–12). Pathological hypoxia may occur due to reduced uterine artery blood flow (13) or an imbalance between in placental oxidative stress and antioxidant defense mechanisms (14), with direct impact on fetal growth and development.

Beyond its role in oxygen and nutrient exchange, the placenta actively communicates with the maternal system. Under hypoxic conditions, it secretes proinflammatory cytokines, immunostimulatory factors, and vasoactive mediators into the maternal circulation (15), thereby influencing maternal physiology and pregnancy adaptations. Among these molecular signatures, circulating cell-free mitochondrial DNA (ccf-mtDNA) has emerged as a potential biomarker of cellular stress and immune activation (16). We recently showed that trophoblast cells release mtDNA fragments in response to oxidative stress and that these fragments circulate in maternal blood as free (“naked”) DNA or enclosed within extracellular vesicles (EVs) (16, 17).

Gestational chronic intermittent hypoxia (CIH), a pattern of fluctuating oxygenation characteristic of obstructive sleep apnea, has emerged as a clinically relevant experimental model of gestational hypoxia or gestational sleep apnea (13). In mice, exposure to gestational CIH induces preeclampsia-like features and fetal growth restriction (18), and when occurring in late gestation, results in long-term adverse neurobehavioral outcomes in offspring (19, 20).

However, the effects of intermittent hypoxia, such as that experienced during gestational sleep apnea, on maternal inflammation and placental stress responses are not fully understood. Furthermore, whether CIH alters ccf-mtDNA levels and whether such changes are associated with placental stress and impaired placental growth remain unknown.

Therefore, the objective of this study was to investigate the effects of gestational CIH on maternal inflammatory responses and placental stress. We hypothesized that gestational CIH would 1) increase proinflammatory cytokines and ccf-mtDNA in maternal circulation, 2) impair placental growth and efficiency, and 3) induce a placental cell stress response characterized by an imbalance between proinflammatory and cell death pathways and the activation of protective mechanisms, such as antioxidant enzymes and autophagy.

## METHODS

### Chemicals and Reagents

Details of chemicals and reagents used in this study are provided in Supplementary Materials (Table S1).

### Animals

Animal experiments were conducted in agreement with the Guide for the Care and Use of Laboratory Animals of the National Institutes of Health and the ARRIVE guidelines. The following animal protocols were approved by the Institutional Animal Care and Use Committee of the University of North Texas Health Science Center (IACUC-2021-041).

All experiments were conducted using Long-Evans timed-pregnant rats (Charles River Laboratories). Rats arrived at the animal facilities on GD6 (GD1=day when vaginal plug is observed; term=22-23 days, body weight on arrival=180-260 g), were single-housed in a 12-hour:12-hour reverse light/dark cycle with lights on at 1900 hour and were allowed to habituate to housing conditions for 8 days prior to any treatment. Food and water were provided *ad libitum*. In total, 19 female rats were purchased as timed-pregnant. At euthanasia, two rats were determined as non-pregnant and removed from downstream analyses. Thus, data from 17 rats were included in this study.

### Experimental Design

To examine the impact of hypoxia during late gestation, timed-pregnant rats received either CIH (n=8) or Normoxia (room air; n=9) starting at 2100 hour for 8 hours during their sleep phase of the circadian cycle on GD15–20.

To induce CIH, the home cages of the pregnant rats were placed into Oxycycler chambers (76.2 × 50.8 × 50.8 cm, BioSpherix, Parish, NY, USA). Animals were allowed to acclimatize to the chambers under normoxic conditions for a period of 6 days prior to starting the CIH protocol, in which oxygen was reduced from 21% (room air) to 10% and then returned to 21% using a 6-minute cycle (10 cycles/hour) over 8 hours/day for a period of 5 days, as we previously described (19, 20). Maternal weight, food intake, and water intake were monitored daily throughout the duration of the treatment. Weights were collected in the morning at the end of the sleep cycle (0800 hour).

### Tissue Sample Collection and Preparation

Rats were euthanized on GD20 during the first 2-5 hours after the last CIH exposure (wake phase). Each animal was anesthetized with isoflurane and euthanized by decapitation, following methods previously described (20–24). Maternal trunk blood was collected in EDTA-coated tubes for plasma isolation, heparinized tubes for whole blood gas analyses, and uncoated tubes for serum isolation. Whole blood collected in uncoated tubes was allowed to clot at room temperature for 30-60 minutes prior to centrifugation for serum collection. Plasma and serum collection tubes were centrifuged at 2000×*g* for 10 minutes at 4°C to isolate plasma (circulating cytokine and markers of oxidative stress analyses) or serum (extracellular vesicle (EV) isolation and characterization, as well as mtDNA quantification). Maternal hematocrit and hemoglobin were assessed as hematologic markers of maternal hypoxia (25). Maternal heparinized whole blood was analyzed for blood gas levels using an automated blood gas analyzer and CO-oximeter system (GEM 3000, Instrumentation Laboratories, Bedford MA, USA).

Fetal hematocrit was measured in one fetus per litter immediately after euthanasia. Fetal trunk blood was collected in heparinized capillary tubes (Fisher Scientific, Waltham MA, USA) that were centrifuged for 3 minutes using a fixed speed Micro-Hematocrit II Centrifuge (Clay Adams, Becton Dickinson, Franklin Lakes NJ, USA). Centrifuged microhematocrit tubes were read on a Micro-hematocrit capillary tube reader (Damon IEC Division, USA) to determine fetal hematocrit. Because accurate hematocrit determination requires immediate sampling, we measured it in a single, randomly selected fetus per litter to avoid delays while reserving the remaining fetuses and placentas for weight measurements. 1-2 placentas/litter and corresponding decidual tissues were randomly selected for rapid flash-freezing in liquid nitrogen to preserve protein and RNA integrity. Flash-frozen placenta/decidua pairs were stored at −80°C for biochemical analyses.

### Fetoplacental Biometrics

Fetal and placental weights (g) were recorded for each pup within litters, and averages were calculated for each litter. Fetal biometrics, including crown-to-rump length (cm) and abdominal girth (cm), were measured for 5-6 pups per litter, and the averages were calculated for each litter.

### Circulating Markers of Oxidative Stress

Circulating oxidized proteins in plasma were measured using the OxiSelect Advanced Oxidative Protein Products (AOPP) Assay Kit with modifications to the manufacturer’s instructions, as previously described (26, 27). Briefly, plasma samples were diluted 1:2 in assay buffer and analyzed in the same plate with and without the reaction initiator to account for plasma background readings. The background values were subtracted from the respective sample readings, and the concentration of AOPP (μM) was quantified using a 4-parameter logistic standard curve.

### Plasma Cytokine Analysis

Plasma cytokines were quantified with MILLIPLEX® Rat Cytokine/Chemokine Magnetic Bead Panel, following a previously described protocol (26, 27). The panel was customized to detect specific cytokines, including (interleukin (IL)-18, IL-1α, IL-1β, IL-6, IL-10, and tumor necrosis factor (TNF)-α). Plasma samples were diluted 1:2 in assay buffer and processed according to the manufacturer’s instructions. All samples, standards, and quality controls were plated in duplicate. Cytokine concentrations were measured using a Luminex® 200 instrument using xPONENT® software version 4.3 (Luminex Corporation, Austin, TX). Quality control values for each cytokine were within the ranges specified by the manufacturer.

### Circulating Cell-free Mitochondrial DNA Quantification

#### DNA Extraction

DNA was extracted from maternal serum samples in two forms: 1) circulating-cell free DNA and 2) DNA encapsulated within isolated EVs. *Circulating-Cell Free DNA:* Circulating-cell free DNA was extracted using the Mag-Bind® Blood & Tissue DNA HDQ extraction kit (Omega Bio-Tek, Norcross, Georgia, USA) following the manufacturer’s specifications for the Mag-Bind® Blood protocol, with modifications as published previously (16, 17). *Extracellular Vesicle DNA:* DNA was extracted from 200 μL of the resuspended isolated EVs isolated from serum samples using the DNeasy Blood & Tissue Kit. The extraction followed the “Purification of Total DNA from Animal Blood or Cells (“Spin-Column Protocol” provided by the manufacturer) and as described in prior publications (16).

#### DNA Quantification

Nuclear DNA and mtDNA were quantified using relative and absolute polymerase chain reaction (PCR) quantification protocols based on Nicklas et al. (28) with some modifications, including the omission of the mitochondrial deletion target. TaqMan chemistry-based approaches were employed for both relative and absolute quantifications. mtDNA was quantified via the mitochondrial D-loop and nuclear DNA was quantified via the β-actin gene.

Relative qPCR was used to assess mtDNA contained in EVs (EV-mtDNA), while absolute qPCR was utilized to measure ccf-mtDNA. Measurements of ccf-mtDNA refer to membrane-bound mtDNA quantities, including but not limited to mtDNA contained in EVs. The qPCR master mix was prepared as follows: 2 μL of each primer (D-loop, 0.625 μM; β-actin, 2.5 μM); 1 μL of each probe (D-loop, 2.5 µM; β-actin, 2.5 µM); 13 μL of TaqMan^TM^ Universal Master Mix II, no UNG; 2 μL of DNA extract. The target sequences for this analysis were:

- Mitochondrial D-loop. Forward: GGTTCTTACTTCAGGGCCATCA; Reverse: GATTAGACCCGTTACCATCGAGAT; Probe: 6FAM-TTGGTTCATCGTCCATACGTTCCCCTTA-TAMRA, (GenBank accession no. X14848)
- Nuclear target β-actin. Forward: GGGATGTTTGCTCCAACCAA; Reverse: GCGCTTTTGACTCAAGGATTTAA; Probe: VIC-CGGTCGCCTTCACCGTTCCAGTT-TAMRA, (GenBank accession no. V01217

For relative qPCR of EV-mtDNA, β-actin was chosen for normalization based on its stable expression across all samples. Absolute quantification of the ccf-mtDNA target was performed using a standard curve of known mtDNA copy number along with each batch of the unknown samples (Sequence: GGTTCTTACTTCAGGGCCATCAATTGGTTCATCGTCCATACGTTCCCCTTAAATAAGACATCTCGATGGTAACGGGTCTAATC). Each plate included the standard curve to interpolate target copy numbers, with R^2^ values and amplification efficiencies assessed to ensure repeatability and to detect potential batch effects. Negative controls were included on every run to monitor off-target mtDNA amplification and ensure the specificity of the assay.

### Extracellular Vesicle Isolation and Characterization

EVs were isolated from 200 μl of maternal serum samples using the ExoQuick^®^ Exosome Precipitation Solution according to the manufacturer’s instructions and as we previously described (16) with some modifications. Briefly, 45 µl of ExoQuick® solution was added to 180 μl of clarified serum and incubated at 4°C for 1 hour. EVs were then pelleted by centrifugation at 1500x*g* for 30 minutes, after which the supernatant was removed. The EV pellet was resuspended in 200 μl of phosphate buffered saline (PBS) and stored at −80°C until further analysis, including DNA extraction or EV characterization. EV characterization was performed by Systems Biosciences using an Exo-Check^TM^ exosome antibody array and nanoparticle tracking analysis (NTA; Nanosight, Version 2.3, Build 2.3.5.0033.7-Beta-7, cat# CSNANO100A-1), as previously described (16). The NTA analysis was performed by Systems Biosciences using a proprietary fluorescence dye, ExoGlow-NTA, which selectively binds to the intact membranes of EVs to enable detection of their size and concentration. Representative images from the Exo-Check^TM^ analysis and nanoparticle tracking analysis are shown in Supplementary Materials (Figure S1).

### Sex Determination of Rat Placentas

Sex was determined in the placentas used for biochemical analyses. Specifically, one placenta was randomly selected from each litter and used for both protein (Western blot) and mRNA expression analyses. DNA from these placentas was extracted using the Extract-N-Amp ^TM^ Tissue PCR kit according to the manufacturer’s instructions. Briefly, placental samples (2-10 mg) were incubated in extraction and tissue preparation solution for 10 minutes, followed by heating at 95 °C for 3 minutes. The reaction was stopped with neutralization solution buffer. A single-step PCR using three primers (Supplementary Materials, Table S2) was performed to determine genetic sex of placental tissues as described by Dhakal et al. (29). The PCR reaction included: 1 μl of DNA, 12 μl of Taq master mix, and 1 μl of primer mix. PCR amplification was carried out on a thermocycler (T100 Thermal cycler, BioRad, Foster City, California, USA) with the following cycling parameters: 94°C for 5 minutes, 35 cycles of denaturing at 94°C for 30 seconds, primer annealing at 55°C for 30 seconds, and extension at 72°C for 42 seconds; 2 cycles at 72°C for 7 minutes. Amplified PCR products were loaded in a 1% agarose gel and were resolved by electrophoresis at 110 Volts for 45 minutes. PCR products from tail DNA of adult male and female rats served as positive controls.

Our analysis revealed that all but one of the randomly selected placentas corresponded to male fetuses (Supplementary Materials (Figure S2). The female placenta was from the CIH group. All placental samples were included in downstream analyses. For transparency, data from the single female placenta are represented using a distinct symbol in all graphs.

### RNA Extraction from Placental and Decidual Tissues

RNA was extracted from one placenta/litter and its corresponding decidua. Tissues (∼100 mg) were homogenized by grinding samples in liquid nitrogen, followed by suspension in QIAzol^®^ Lysis Reagent. RNA extraction was performed using the miRNeasy Mini Kit according to the manufacturer’s protocol. The purity and concentration of the isolated RNA were assessed using a Nanodrop spectrophotometer (NanoDrop One Spectrophotometer, Thermo Scientific, Waltham, MA, USA). All samples exhibited 260/280 absorbance ratios ranging from 1.98 to 2.07, indicating high RNA purity. RNA was diluted to a concentration of 2 μg/μl using RNase-free water.

### cDNA Synthesis and Quantitative Real-Time PCR

RNA was reverse-transcribed into cDNA using Sensiscript RT Kit reagents with RiboGuard RNase inhibitor and oligo-dT primers as previously published (30, 31). cDNA synthesis was carried out on a thermocycler (T100 Thermal cycler, BioRad, Foster City, California, USA) set at 37 °C for 1 hour. cDNA was stored at −20 °C until further analysis.

The expression levels of antioxidant enzymes *(sod1, sod2, catalase)*, cytokines (*tnfα, il-1β, il-6, il-10, il-18),* and the reference gene *gapdh* were measured using qRT-PCR. *Gapdh* was selected as the reference gene for normalization based on its stable expression across all samples. Primer sequences were purchased from Integrated DNA Technologies (IDT, Redwood, CA, USA) and are provided in Supplementary Materials (Table S3).

qRT-PCR was performed as previously described (30, 31). Briefly, the reaction mix consisted of 7.5 μl iQ SYBR Green Supermix; 1.2 μl primer mix (prepared with 50 μl of 100 mM forward primer, 50 μl of 100 mM reverse primer, and 100 μl RNase-free water); 1.8 μl cDNA; 4.5 μl RNase-free water. Non-template controls were included for each primer set in duplicate for both target and reference genes. PCR reactions were performed using a CFX96 Real-Time PCR Detection System with CFX Maestro Software v.2.3 (Bio-Rad, Foster City, CA, USA), with the following cycling parameters: initial denaturation at 95 °C for 3 minutes, followed by 40 cycles of 95 °C for 10 seconds, and 60 °C for 1 minute. A melt curve was generated at the end of the cycling protocol to verify primer specificity. Gene expression was analyzed using the ι1Ct, and results are expressed as 2^-ΔΔCt^.

### Western Blot

Protein was extracted from one placenta/litter and its corresponding decidua using a T-PER tissue protein extraction reagent supplemented with protease inhibitor cocktail tablets. Protein concentrations were determined for each sample using the Pierce^TM^ bicinchoninic acid (BSA) protein assay. Samples were denatured with β-mercaptoethanol and heated to 99°C for 5 minutes. Equal amounts of protein (15-30 μg) were loaded onto 4-15% of Mini-Protean TGX stain-Free Precast gels (Bio-Rad, Foster City CA, USA) and electrophoresed for 2 hours at 100 Volts. Proteins were transferred to nitrocellulose membranes using a Trans-Blot Turbo Transfer System (BioRad, Foster City, CA, USA) for 10 minutes. Membranes were then blocked for 1 hour with 3% Bovine Serum Albumin (BSA) or 5% non-fat milk in tris-buffered saline with Tween-20 (T-BST). Primary antibodies were diluted in 3% BSA or 5% non-fat milk in T-BST and incubated overnight at 4°C. After washing with T-BST, membranes were incubated with the appropriate secondary antibody for 1 hour at room temperature. Data acquisition was performed using a) Odyssey CLx LI-COR (LI-COR Biosciences), and analysis was conducted using Image Studio v. 5.2 (LI-COR Biosciences) or b) AZURE 300 (Biosystems), and analysis was conducted using ImageJ software (version 13.0.6, NIH, USA). Protein signals were normalized to total protein using Ponceau staining, with quantification performed in ImageJ software (version 13.0.6). All immunoblots are presented and details regarding antibody usage and protocols are provided in Supplementary Materials (Figure S3-15 and Table S4, respectively).

### Statistical Analysis

Data distributions were assessed using the Shapiro-Wilk test. Outlier detection was performed using the ROUT (Robust regression and Outlier) method with a coefficient Q=1%. Maternal weights and food intake data were analyzed using two-way ANOVA with repeated measures (or mixed model), applying the Geisser-Greenhouse correction, followed by Sidak’s multiple comparisons test. For other comparisons, unpaired t-tests were used for normally distributed data with equal variances, Welch’s t-tests were applied for data with unequal variances, while Mann-Whitney *U* tests were used for data that were not normally distributed. Data are presented as mean ± standard deviation (SD), unless otherwise indicated. For qPCR data analysis, statistical tests were performed on ι1Ct values, and results are reported as 2^-ΔΔCt^. The significance level was set to α = 0.05. All statistical analyses were conducted in Prism Version 10.4 (GraphPad, San Diego, CA), unless otherwise specified. Exact p values are reported for all analyses.

## RESULTS

### Maternal Outcomes

#### Maternal hematologic adaptive responses to hypoxia

At euthanasia, we assessed maternal hematocrit and hemoglobin, as markers of an adaptive response to hypoxia. Maternal hematocrit (Normoxia (n=9): 27.4% ± 3.40 vs. CIH (n=8): 31.0% ± 3.3, Unpaired t-test, p=0.046) and hemoglobin concentrations (Normoxia (n=9): 10.7 g/dL ± 1.19 vs. CIH (n=7): 12.01 g/dL ± 1.11, Unpaired t-test, p=0.042) were higher in the CIH group compared to Normoxia.

#### Maternal body weights, and food and water intakes

A two-way repeated measures ANOVA (treatment x gestational age) revealed a significant interaction effect on maternal body weight (Geisser-Greenhouse-adjusted F_(5, 75)_=4.93, p=0.002; Figure 1A). The main effect of treatment was not significant (F_(1, 15)_=0.0004, p=0.98), whereas the main effect of gestational age was (Geisser-Greenhouse-adjusted F_(2.38, 35.8)_=209.4, p<0.0001). Within the Normoxia group, maternal body weights increased progressively from GD15 to GD20 (post-hoc Sidák’s multiple comparisons test, all p<0.05). In contrast, body weights in CIH-exposed dams did not change from GD15 to GD16 (p=0.999), progressively increased from GD16 to GD19 (all p<0.05), and did not change from GD19 to GD20 (p=0.318). The cumulative weight gained by normoxic dams was significantly greater than that of CIH-exposed dams from GD15 to GD20 (Normoxia (n=9): 65.7 g ± 13.02 vs. CIH (n=8): 46.20 g ± 14.16, Unpaired t-test, p=0.0099). These patterns demonstrate that normal gestational weight gain was blunted under CIH exposure.

**Figure 1.**
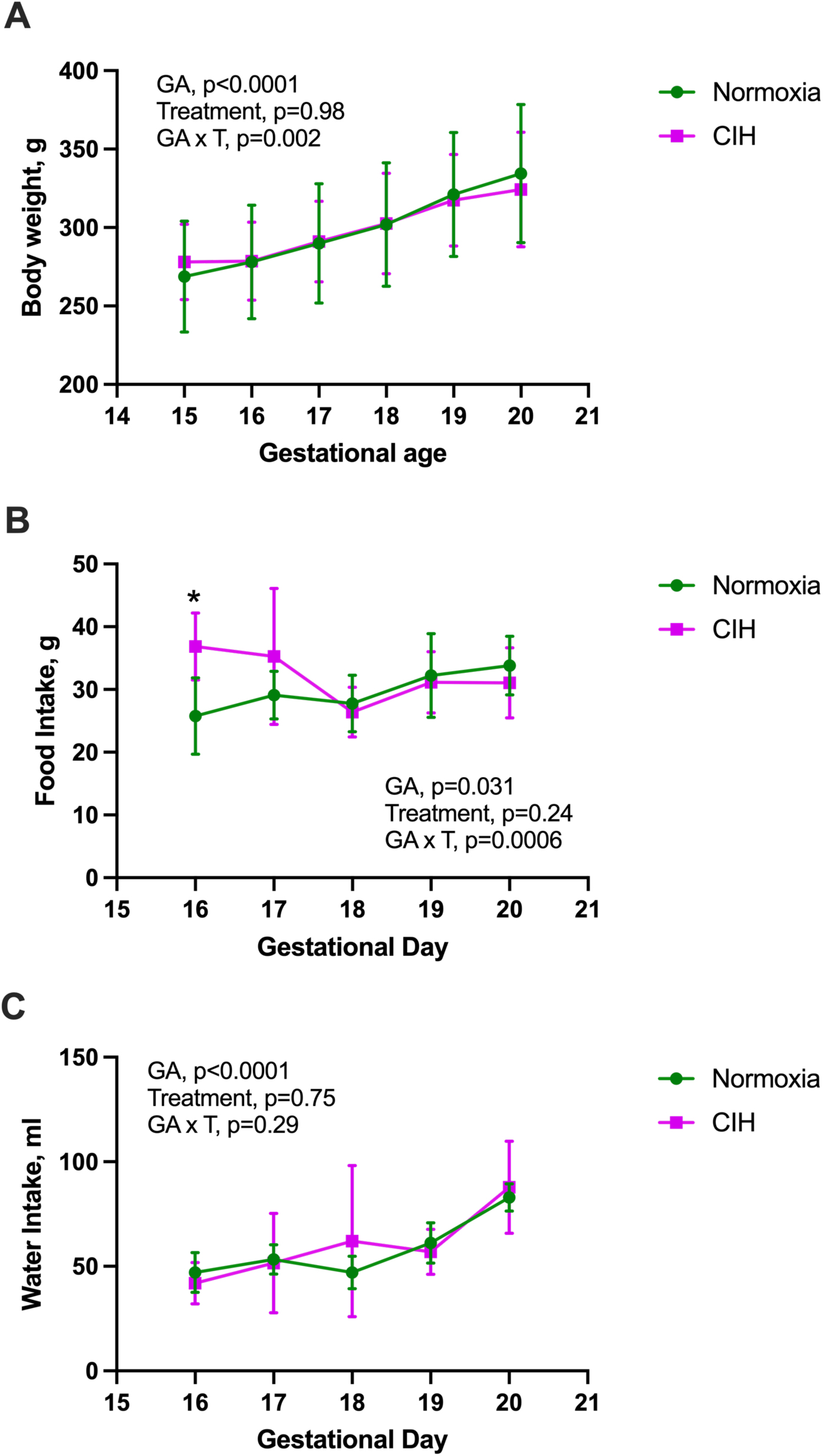
Maternal body weights, and food and water intakes. (A) Changes in maternal body weight with advancing gestational age (GA, day 15-20) differs between chronic intermittent hypoxia (CIH) and Normoxia groups (treatment x gestational age interaction). Two-way ANOVA with repeated measures with Geisser-Greenhouse correction, followed by Sidak’s multiple comparisons test. Normoxia, n=9; CIH, n=8. (B) Food intake was higher in the CIH group on gestational day (GD) 16 but showed no differences thereafter. Mixed-effects model with Geisser-Greenhouse correction, followed by Sidak’s multiple comparisons test. Normoxia, n=8-9; CIH, n=7-8. *p=0.012. (C) There was no significant interaction effect (treatment x gestational age) on maternal water intake. Mixed-effects model with Geisser-Greenhouse correction. Normoxia, n=8-9; CIH, n=7-8. Data are presented as means ± SD.

A mixed-effects model (treatment x gestational age) identified a significant interaction effect on maternal food intake (Geisser-Greenhouse-adjusted F_(4, 58)_=5.73, p=0.0006; Figure 1B). The main effect of treatment was not significant (F_(1, 15)_=1.49, p=0.24), whereas food intake varied across gestation (Geisser-Greenhouse-adjusted F_(2.68, 38.86)_=3.42, p=0.031). Post-hoc Sidák’s multiple comparisons test revealed no significant differences in food intake within each treatment across gestation (all p>0.05). Between groups, the only difference occurred on GD16, when the CIH group consumed more chow compared to the Normoxia group (p=0.012).

There was no significant interaction effect (treatment x gestational age) on maternal water intake (mixed-effects model, Geisser-Greenhouse-adjusted F_(4, 59)_=1.27, p=0.29; Figure 1C).

#### Systemic markers of maternal oxidative stress and inflammation

We measured plasma AOPP levels and concentrations of pro-inflammatory (TNF-α, IL-1β, IL-18) and anti-inflammatory (IL-10) cytokines as markers of systemic oxidative stress and inflammation. Levels of AOPP in maternal plasma did not differ between Normoxia and CIH-exposed dams (Normoxia (n=8): 140.5 μM ± 19.07 vs. CIH (n=8): 115.2 μM ± 37.20, Welch’s t-test, p=0.116). Circulating TNF-α (Unpaired t-test, p=0.482; Figure 2A), IL-6 (Welch’s t-test, p=0.352; Figure 2B), and IL-1β (Welch’s t-test, p=0.222; Figure 2C) were not affected by CIH. However, IL-18 concentrations were higher in the CIH group compared to Normoxia (Unpaired t-test, p=0.047; Figure 2D). There were no group differences in concentrations of IL-10 (Unpaired t-test, p=0.870; Figure 2E).

**Figure 2.**
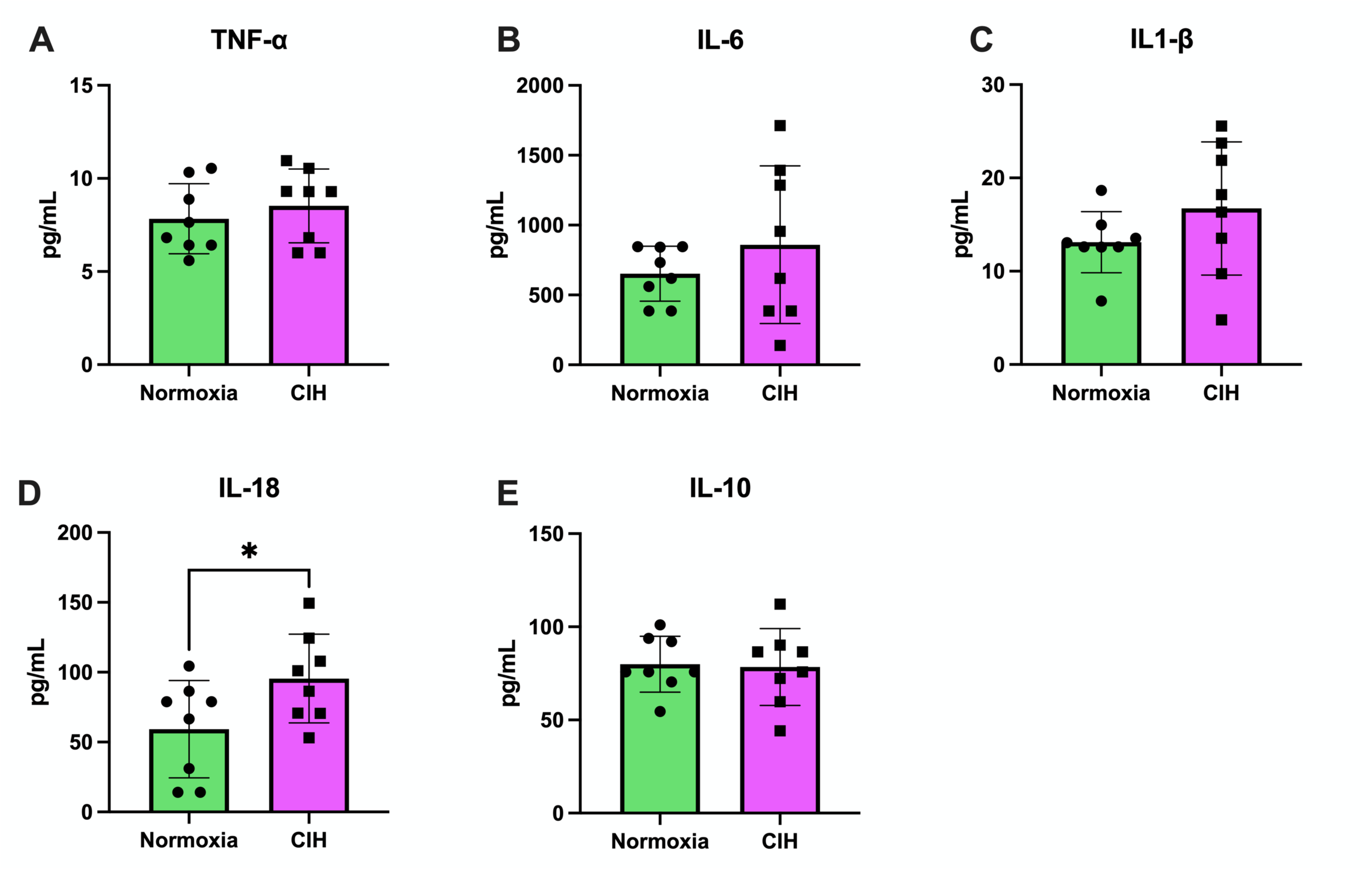
Maternal plasma cytokines. A) Tumor necrosis factor alpha (TNF-α) concentrations in maternal plasma were not different between the chronic intermittent hypoxia (CIH) and Normoxia groups. Unpaired t-test. Normoxia, n=8; CIH, n=8; p>0.05. (B) Interleukin-6 (IL-6) concentrations in maternal plasma were not different between groups. Welch’s t-test. Normoxia, n=8; CIH, n=8; p>0.05. (C) Interleukin-1β (IL-1β) concentrations in maternal plasma were not different between groups. Welch’s t-test. Normoxia, n=8; CIH, n=8; p>0.05. (D) Interleukin-18 (IL-18) concentrations were higher in the CIH group compared to the Normoxia group. Unpaired t-test. Normoxia, n=8; CIH, n=8; p=0.047. (E) Interleukin-10 (IL-10) concentrations in maternal plasma were not different between groups. Unpaired t-test. Normoxia, n=8; CIH, n=8; p>0.05. Data are presented as means ± SD.

Because ccf-mtDNA and EV-mtDNA are emerging markers of cellular stress and inflammation (17, 32–37), we assessed both fractions in maternal serum. ccf-mtDNA (Unpaired t-test, p=0.627, Figure 3A) and EV-mtDNA (Mann-Whitney *U* test, p=0.67, Figure 3B) were not affected by CIH exposure. Isolated EVs had exosome-like traits including exosome-associated protein marker expression and size (Supplementary Materials, Figure S1). EV size was similar between groups (Welch’s t-test, p=0.367, Figure 3C) but EV concentrations were lower in the CIH group (Unpaired t-test, p=0.011, Figure 3D), suggesting greater mtDNA content per EV.

**Figure 3.**
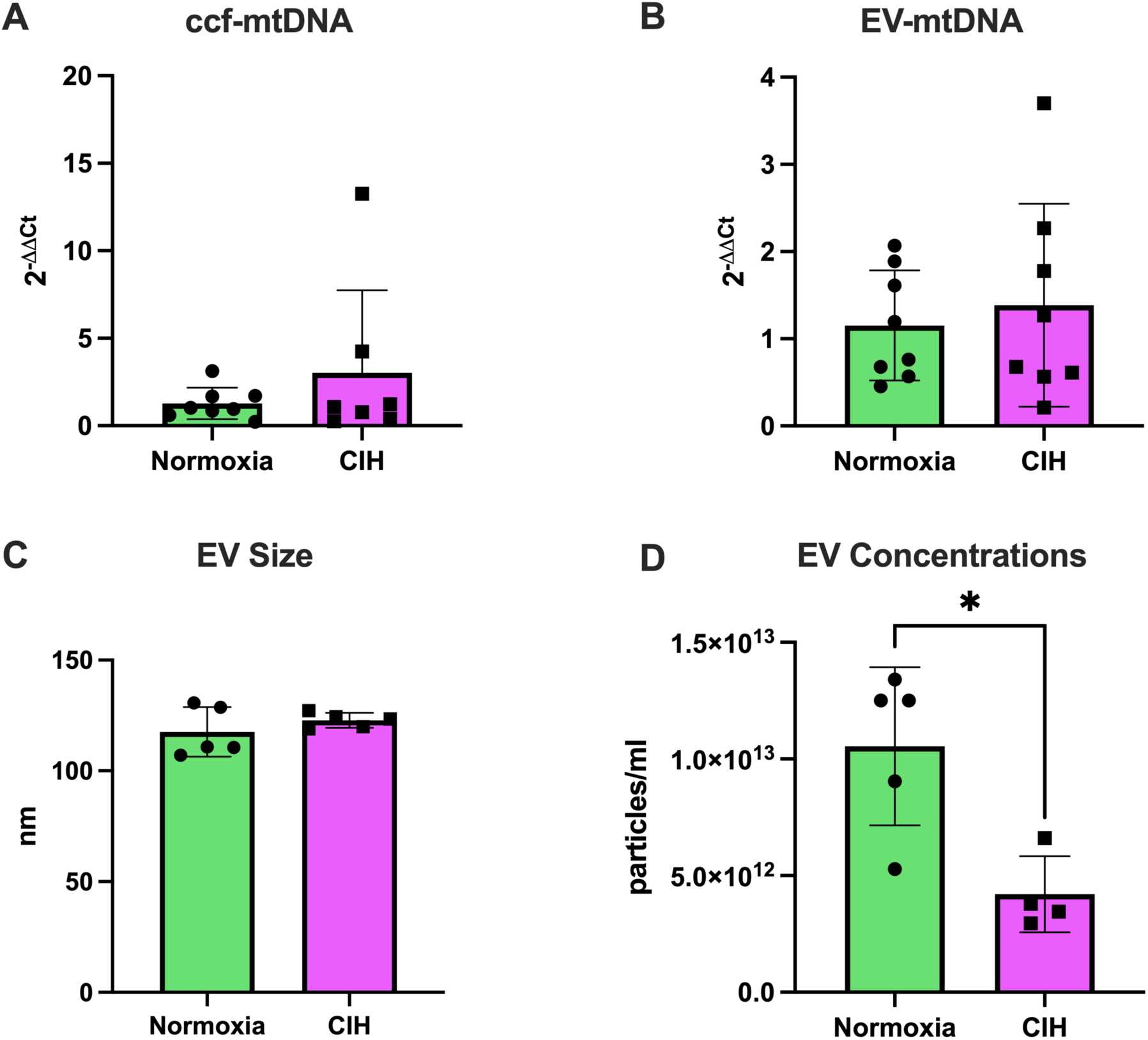
Circulating cell-free mitochondrial DNA and extracellular vesicle characteristics. (A) Circulating cell-free mitochondrial DNA (ccf-mtDNA) levels were not different between the chronic intermittent hypoxia (CIH) and Normoxia groups. Unpaired t-test. Normoxia, n=8; CIH, n=8; p>0.05. (B) Extracellular vesicle (EV)-contained mitochondrial DNA (EV-mtDNA) content did not differ between groups. Unpaired t-test. Normoxia, n=5; CIH, n=5; p>0.05. (C) Size of EVs in maternal serum was not different between groups. Welch’s t-test. Normoxia, n=5; CIH, n=5; p>0.05. (D) Concentrations of EVs in maternal serum were lower in the CIH group compared to the Normoxia group. Unpaired t-test. Normoxia, n=5; CIH, n=4; *p=0.011. Statistics for mtDNA levels were performed on ΔCt values and are presented as 2^-ΔΔCt^. Data are presented as means ± SD.

### Perinatal Outcomes

#### Fetal hypoxia and fetoplacental biometrics

We first determined whether CIH affected fetal oxygen-carrying capacity and adaptive responses via measurements of fetal hematocrit. CIH did not affect fetal hematocrit levels (Normoxia (n=9): 31.5% ± 6.63 vs. CIH (n=6): 32.5% ± 10.5, Mann-Whitney *U* test, p=0.509). Furthermore, CIH did not impact litter size (Normoxia (9): 11.9 ± 2.7 pups vs. CIH (8): 10.5 ± 2.3 pups, Unpaired t-test, p=0.278). Fetal body weights (Unpaired t-test, p=0.486; Figure 4A), abdominal girth (Mann-Whitney *U* test, p=0.864, Figure 4B), and crown-to-rump length (Welch’s t-test, p=0.372, Figure 4C) did not differ between groups. Placental weights were higher in the CIH group compared to Normoxia (Mann-Whitney *U* test, p=0.015, Figure 4D), while fetal-to-placental weight ratios were lower in the CIH group (Unpaired t-test, p=0.0006, Figure 4E). We observed resorptions in two dams exposed to CIH (Dam 1: resorptions=4, litter size=8; Dam 2: resorptions=1, litter size=11). There were no resorptions in the Normoxia group. Collectively, these findings indicate that CIH exposure during rat pregnancy increases placental size with no effects on fetal growth, suggesting a reduction in placental efficiency.

**Figure 4.**
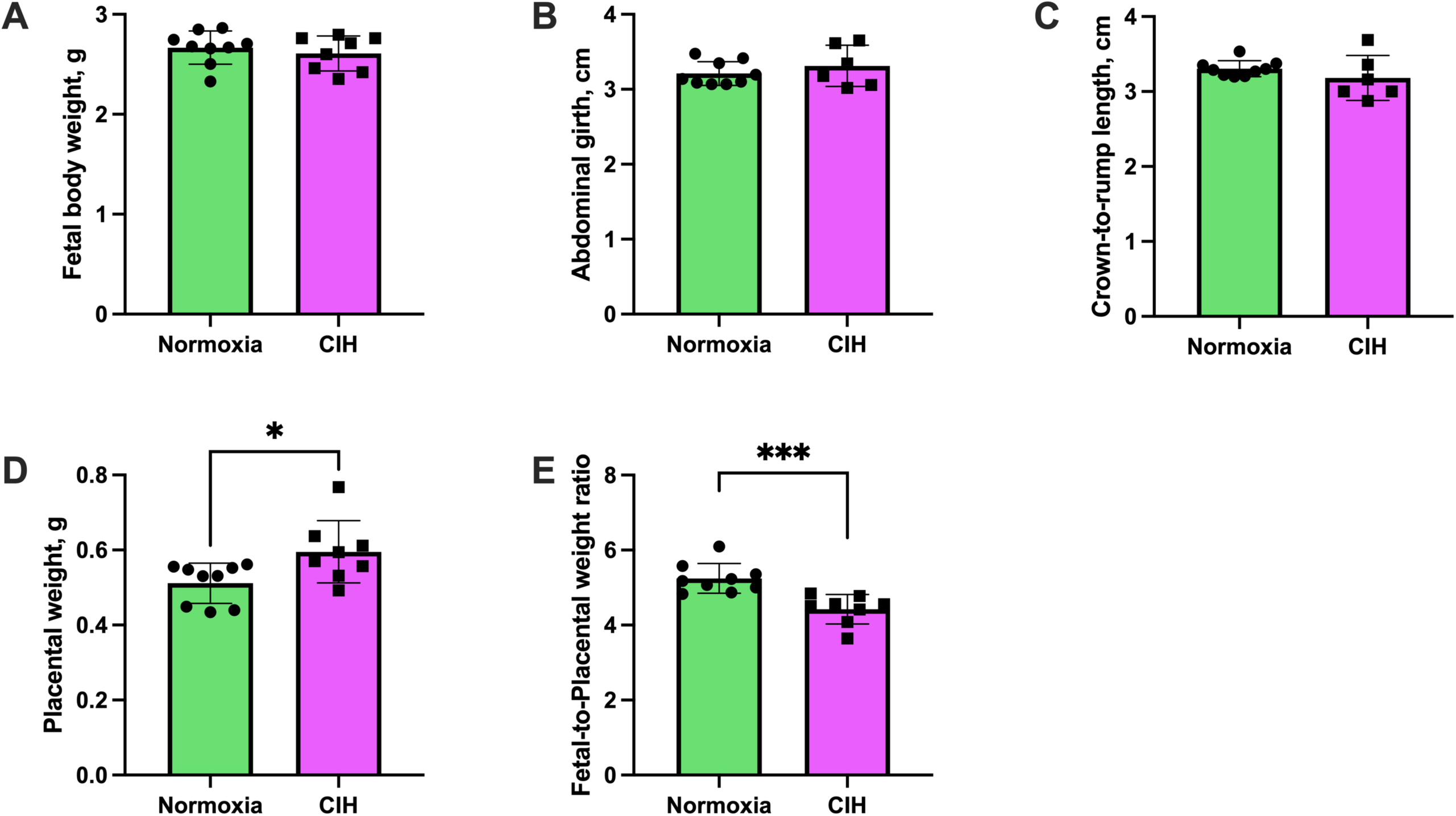
Fetoplacental biometrics. (A) Fetal body weight was not affected by exposure to chronic intermittent hypoxia (CIH). Unpaired t-test. Normoxia, n=9; CIH, n=8; p>0.05. (B) Abdominal girth was not altered by CIH exposure. Mann-Whitney *U*-test. Normoxia, n=6; CIH, n=6; p>0.05. (C) Crown-to-rump length did not differ between groups. Welch’s t-test. Normoxia, n=9; CIH, n=6; p>0.05. (D) Placental weights were higher in the CIH group compared to Normoxia. Mann-Whitney *U*-test. Normoxia, n=9; CIH, n=8. *p=0.015. (E) Fetal-to-placental weight ratios were reduced in the CIH group. Unpaired t-test. Normoxia, n=9; CIH, n=8. ***p=0.0006. For panels (A, D, E), each data point represents the mean values of all pups from a single dam. For panels (D, E), each data point represents the mean values of 2–3 pups sampled from the left and right uterine horns of each dam. Data are presented as means ± SD.

#### Placental Stress Response Mechanisms

Because we observed alterations in placental growth and efficiency (fetal-to-placental weight ratios), we next assessed the placental stress response to CIH, which we defined as an imbalance between the activation of proinflammatory, cell stress, and cell death pathways, and the activation of cellular defense mechanisms, such as antioxidant enzymes and autophagy.

#### Placental inflammation, cellular stress responses, and cell death pathways

To examine placental inflammation, we measured mRNA expression of proinflammatory (*tnfα, il-6*, *il-1β, il-18)* and anti-inflammatory cytokines (*il-10).* There were no differences between groups in placental mRNA expression of *tnfα* (Unpaired t-test, p=0.211, Figure 5A), *il-6* (Welch’s t-test, p=0.724, Figure 5B), *il-1β* (Mann-Whitney *U* test, p=0.072; Figure 5C), *il-18* (Unpaired t-test, p=0.523, Figure 5D), and *il-10* (Unpaired t-test, p=0.524, Figure 5E).

**Figure 5.**
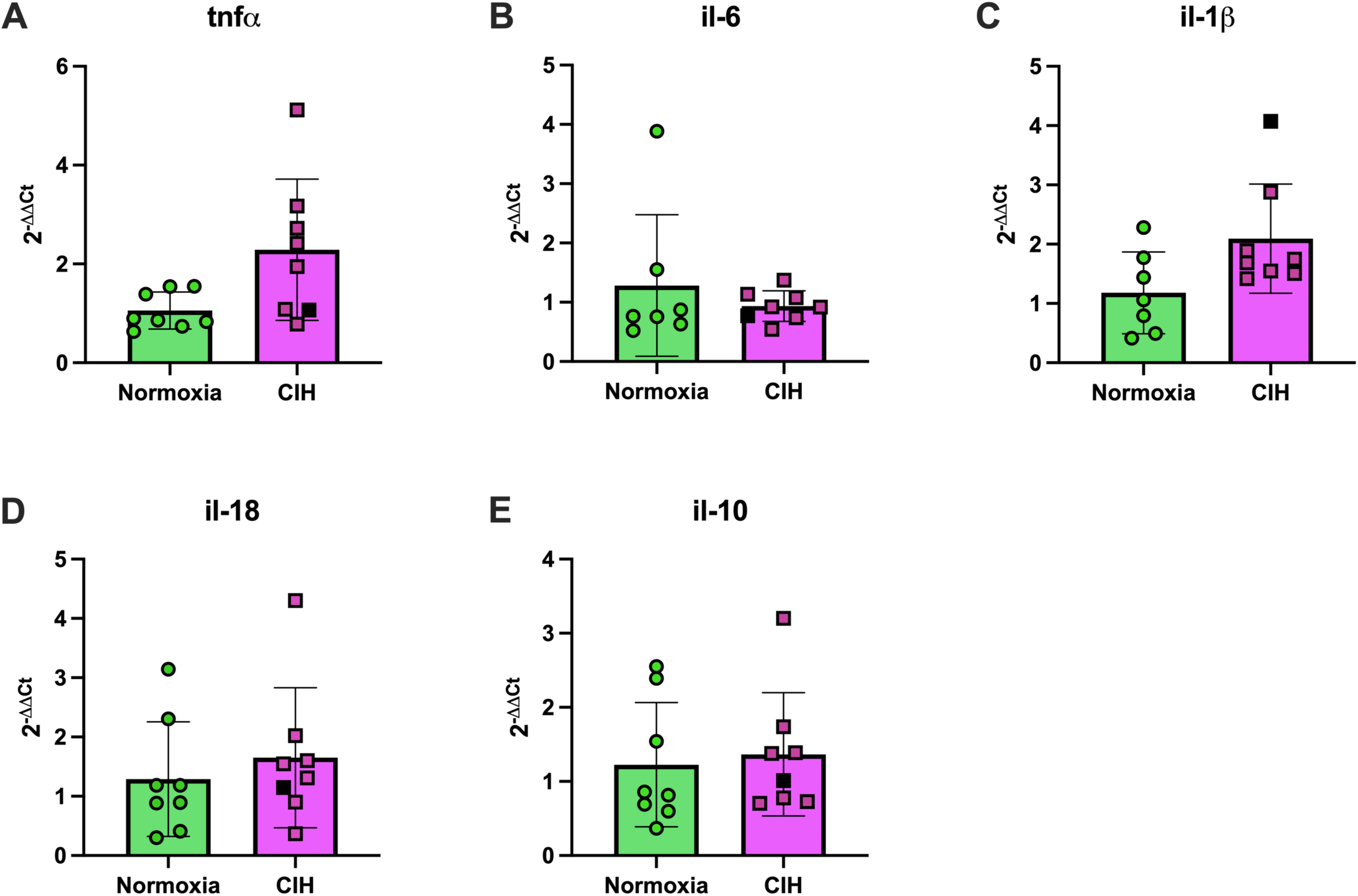
Placental cytokine mRNA expression. Relative mRNA expression levels (2^-ΔΔCt^) of (A) tumor necrosis factor alpha (*tnf-α*), (B) interleukin-6 (*il-6*), (C) interleukin-1β (*il-1β*), (D) interleukin-18 (*il-18*), and (E) interleukin-10 (*il-10*) in placental tissues from rats exposed to chronic intermittent hypoxia (CIH) or Normoxia during gestation (gestational days 15-20). No differences were observed between groups for any cytokine, p>0.05. Unpaired t-tests were used for normally distributed data with equal variances (*tnf-α*, *il-18*, *il-10*; Normoxia, n=8; CIH, n=8), and Welch’s t-tests were used for data with unequal variances (*il-6*; Normoxia, n=7; CIH, n=8). Mann-Whitney *U*-test was used for non-normally distributed data (*il-1β*; Normoxia, n=7; CIH, n=8). Data are presented as means ± SD. Black square symbols indicate data from the single placenta associated with a female fetus; all other data points represent male fetuses.

Activation of mitogen activated protein kinase (MAPK) pathways is a canonical response to cellular stress (38, 39). Therefore, we quantified expression of phosphorylated (activated) and total MAPKs (p38, p44/42). There were no differences in protein expression of phosphorylated and total p38 MAPK (Unpaired t-test, p=0.581 and p=0.538, respectively; Figures 6A-B & Supplementary Materials, Figure S3) and p44/42 MAPK (ERK1/2; Unpaired t-test, p=0.897 and p=0.233, respectively; Figures 6C-D & Supplementary Materials, Figure S4). Representative Western blot images for p38 MAPK and p44/42 (ERK1/2) are shown in Figure 6E and Figure 6F, respectively. Like MAPKs, total caspase-3 and cleaved caspase-3, a terminal effector of apoptosis, were not affected by CIH (Unpaired t-test, p=0.518 and p=0.373, respectively; Figure 7A-B & Supplementary Materials, Figure S5). Representative images of these proteins are illustrated in Figure 7C. Taken together, these data suggest that exposure to CIH in late pregnancy does not induce placental inflammation or cell death.

**Figure 6.**
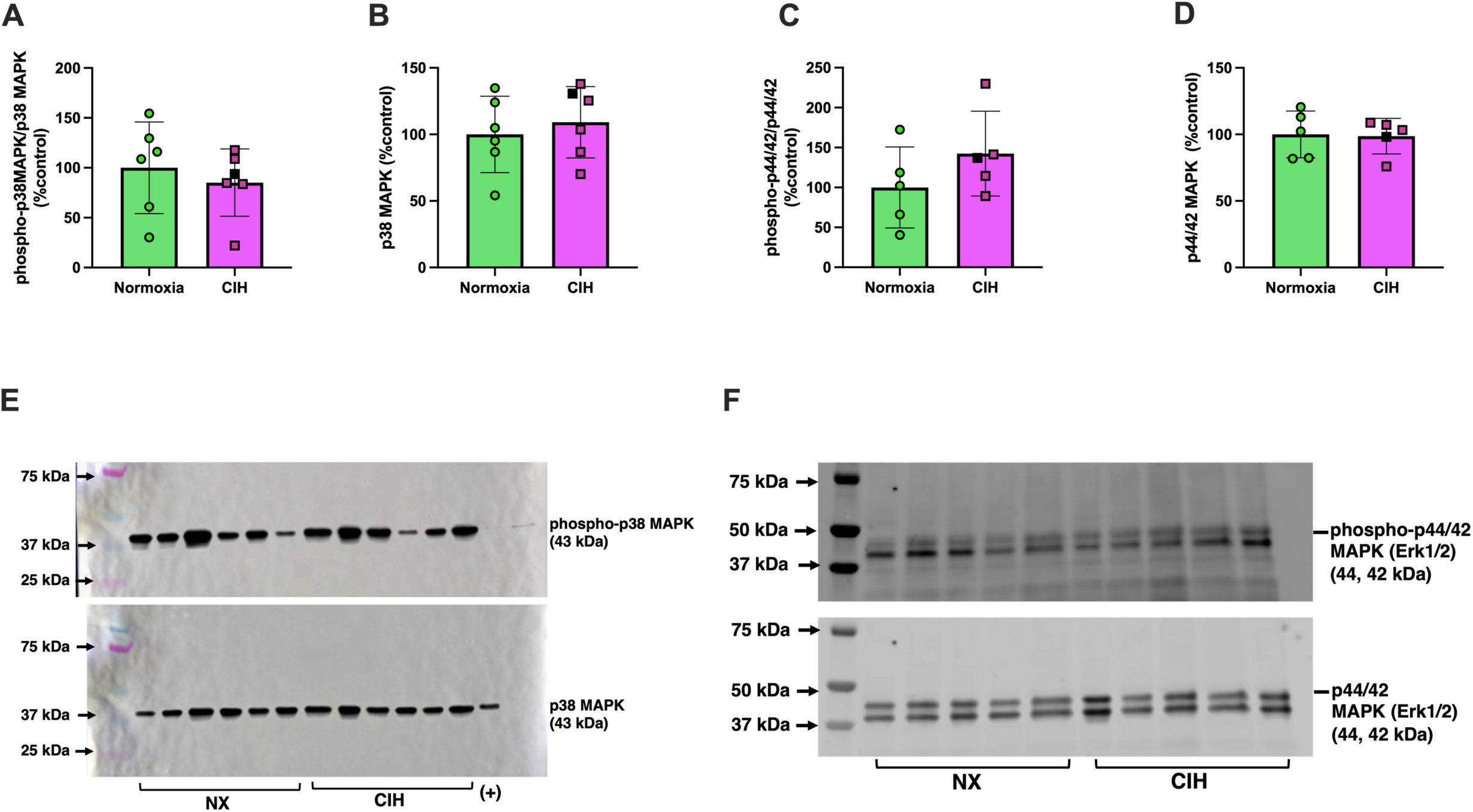
Placental protein expression of placental mitogen-activated protein kinases. Relative expression levels of (A) phosphorylated p38 MAPK, (B) total p38 MAPK, (C) phosphorylated p44/42 (ERK1/2), (D) total p44/42 (ERK1/2) in placental tissues from rats exposed to chronic intermittent hypoxia (CIH) or Normoxia during gestation (gestational days 15-20). (E) and (F) are representative images of Western blot images for p38 MAPK and p44/42 (ERK1/2), respectively. Protein expression levels for phosphorylated forms were calculated as the ratio of phosphorylated levels to total protein, normalized to total loaded protein from Ponceau staining, and expressed as a percentage of Normoxia levels (%control). Total protein levels were normalized similarly. No differences were observed between groups for any protein (p>0.05). Unpaired t-tests were used for normally distributed data with equal variances (p38 MAPK, phospho-p44/42 (ERK1/2), and total p44/42 (ERK1/2); Normoxia, n=5-6; CIH, n=5-6), and Welch’s t-tests were used for data with unequal variances (phospho-p38 MAPK; Normoxia, n=6; CIH, n=6). Data are presented as means ± SD. Black square symbols indicate data from the single placenta associated with a female fetus; all other data points represent male fetuses. MAPK, mitogen-activated protein kinase; ERK1/2, extracellular signal-regulated kinase 1 and 2.

**Figure 7.**
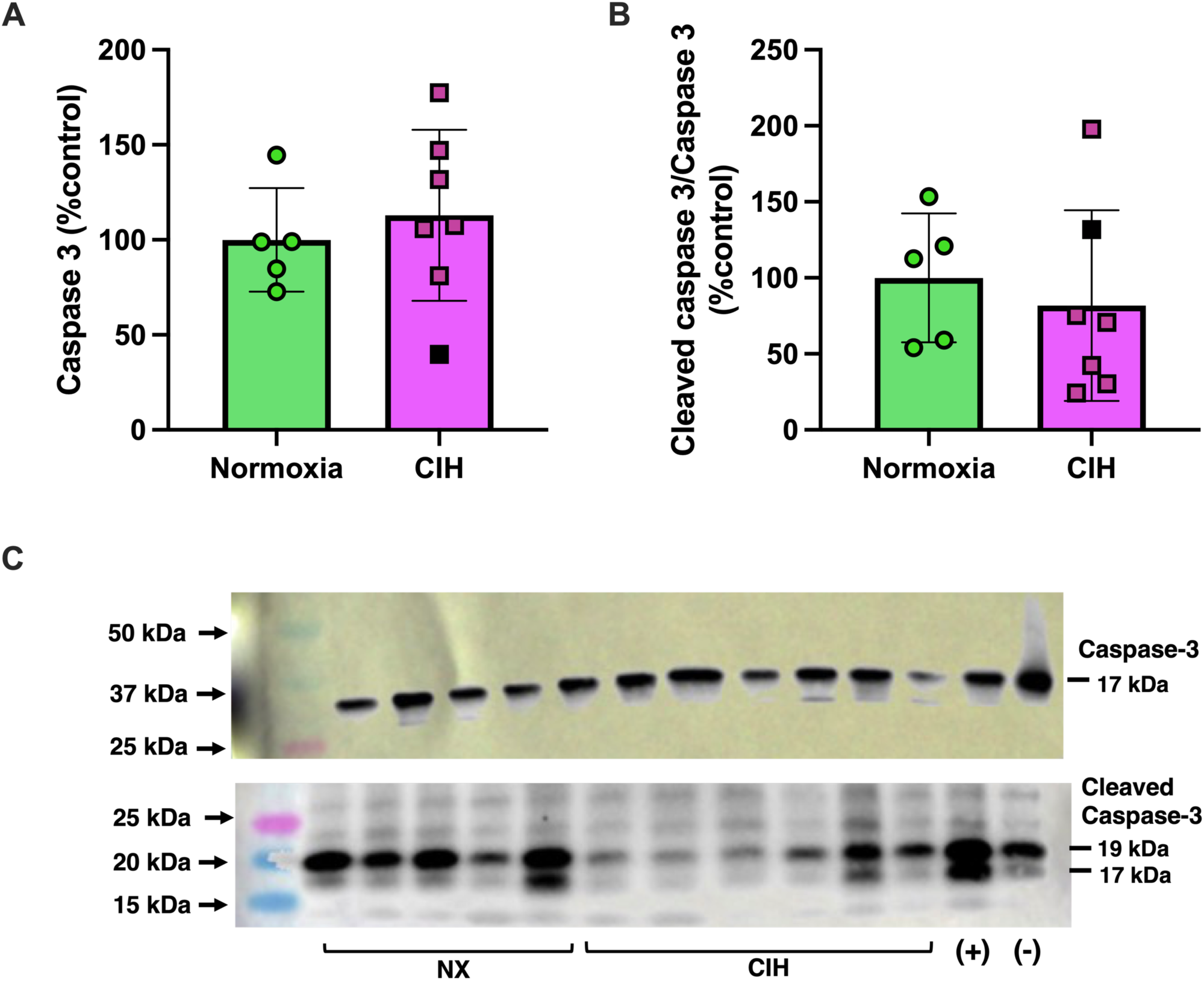
Placental protein expression of placental cleaved caspase-3 and total caspase-**3.** Relative expression levels of (A) caspase-3 and (B) cleaved caspase-3 in placental tissues from rats exposed to chronic intermittent hypoxia (CIH) or Normoxia during gestation (gestational days 15-20). (C) includes representative Western blot images for caspase-3 and cleaved caspase-3. Protein expression levels for cleaved caspase-3 were calculated as the ratio of cleaved caspase-3 to total caspase-3, normalized to total loaded protein from Ponceau staining, and expressed as a percentage of Normoxia levels (%control). Total caspase-3 protein levels were normalized similarly. No differences were observed between groups for any protein (p>0.05). Unpaired t-tests were used for all comparisons. Normoxia, n=5; CIH, n=6. Data are presented as means ± SD. Black square symbols indicate data from the single placenta associated with a female fetus; all other data points represent male fetuses.

#### Placental antioxidant enzymes and autophagy markers

To determine whether CIH stimulates placental antioxidant responses, we quantified expression of the key antioxidant enzymes, Catalase, SOD2, SOD1. Placental mRNA expression of *catalase* (Unpaired t-test, p=0.048; Figure 8A) and *sod2* (Mann-Whitney *U* test, p=0.038; Figure 8B) were higher in the CIH group, while *sod1* did not differ between groups (Unpaired t-test, p=0.555; Figure 8C). CIH did not affect protein content of Catalase (Mann-Whitney *U* test, p=0.965; Figure 8D & Supplementary Materials, Figure S6), SOD2 (Unpaired t-test, p=0.384; Figure 8E & Supplementary Materials, Figure S7), and SOD1 (Unpaired t-test, p=0.503; Figure 8F & Supplementary Materials, Figure S6).

**Figure 8.**
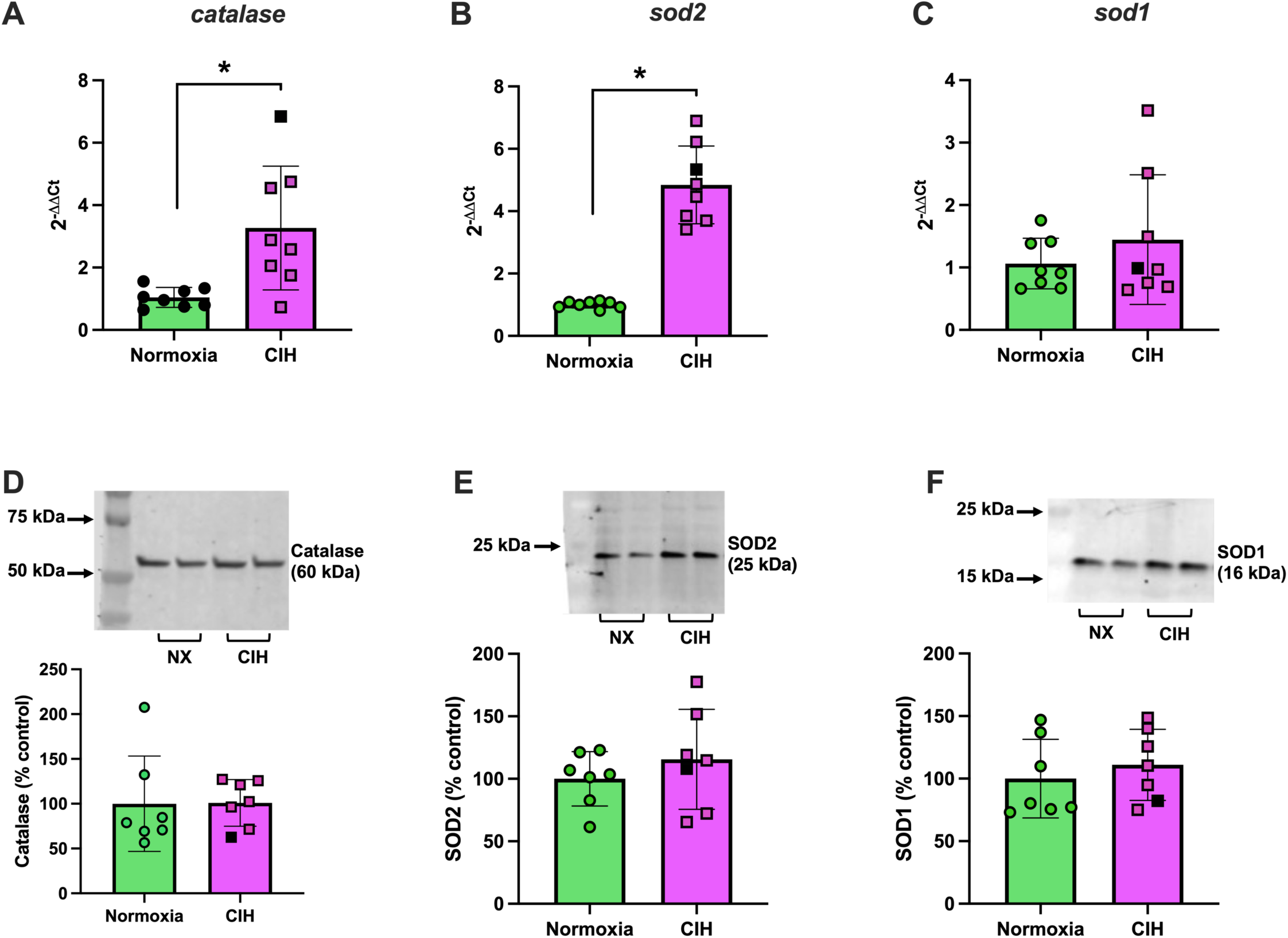
Placental expression of antioxidant genes and proteins. Relative mRNA expression levels (2^-ΔΔCt^) of (A) *catalase*, (B) *sod2*, (C) *sod1*, and protein expression levels of (D) Catalase, (E) SOD2, (F) SOD1 in placental tissues from rats exposed to chronic intermittent hypoxia (CIH) or Normoxia during gestation (gestational days 15-20). Statistics for mRNA expression levels were performed on ΔCt values and are presented as 2^^-ΔΔCt^. Protein expression levels were calculated as the ratio of protein, normalized to total loaded protein from Ponceau staining, and expressed as a percentage of Normoxia levels (%control). mRNA expression of (A) *catalase* (p=0.048) and (B) *sod2* (p=0.038) were higher in CIH (n=8) compared to Normoxia (n=8) group. There were no group differences in *sod1* (n=8/group) mRNA expression and protein levels of Catalase, SOD1, SOD2 (n=7/group), p>0.05. Unpaired t-tests were used for normally distributed data with equal variances (*catalase*, *sod1*, Catalase, SOD1, SOD2), and Mann-Whitney *U* test were used for non-normally distributed data (*sod2*) data. Data are presented as means ± SD. Black square symbols indicate data from the single placenta associated with a female fetus; all other data points represent male fetuses. SOD, superoxide dismutase.

Autophagy is a degradation mechanism that regulates cellular homeostasis; therefore, we assessed Beclin-1 (initiator of autophagy), p62 (marker and regulator of autophagy), and LC3A/B (marker of autophagosome formation) (40). Phosphorylated Beclin-1 (Unpaired t-test, p=0.119, Figure 9A & Supplementary Materials, Figure S8) did not differ between groups, while total Beclin-1 (Mann-Whitney *U* test, p=0.01, Figure 9B & Supplementary Materials, Figure S8) and p62 (Unpaired t-test, p=0.006, Figure 9C & Supplementary Materials, Figure S9) were higher in the CIH compared to Normoxia group. CIH did not change LC3A/B expression (Unpaired t-test, p=0.117, Figure 9D & Supplementary Materials, Figure S10). Representative Western blot images are shown in Figure 9E.

**Figure 9.**
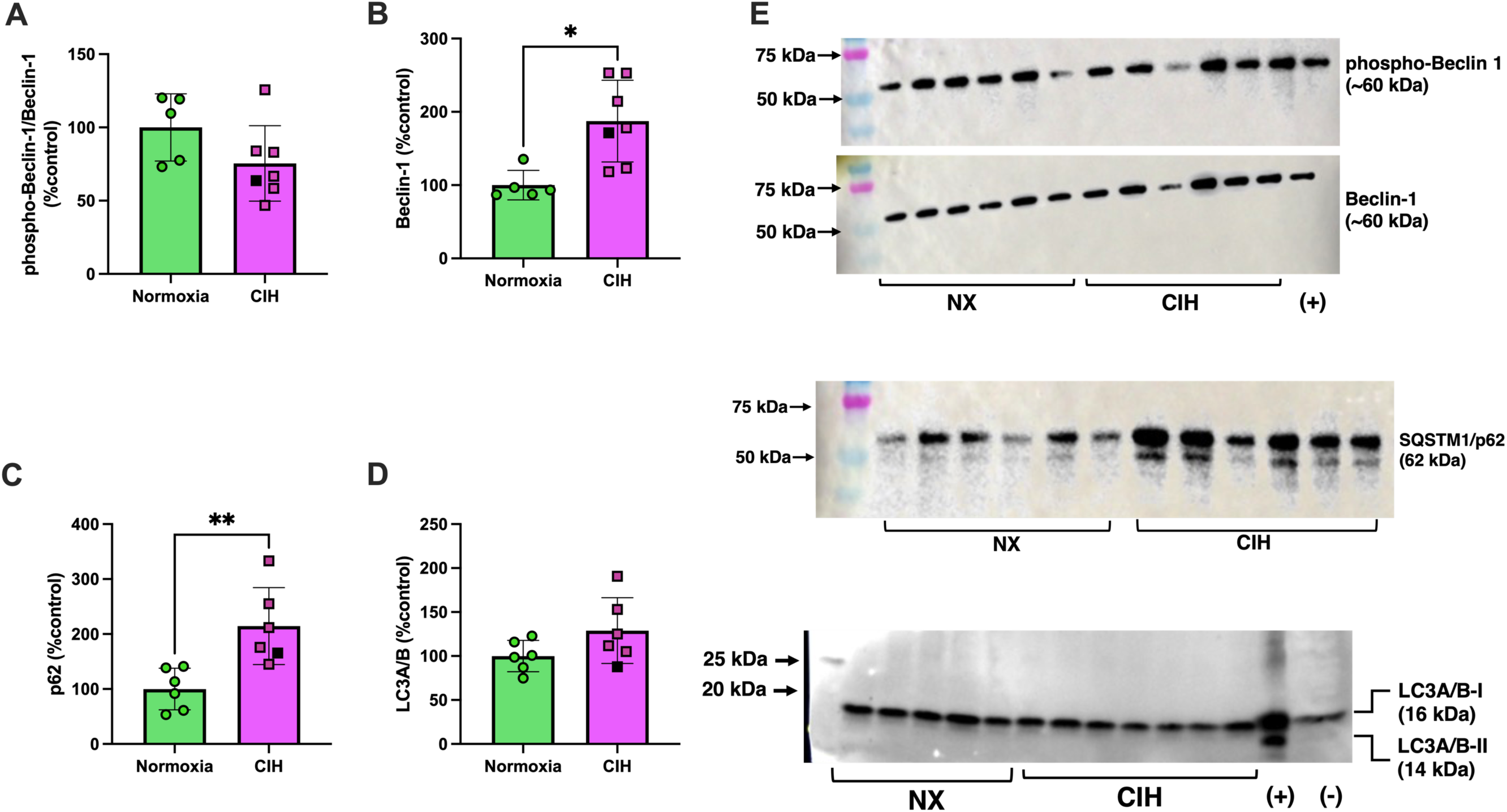
Placental expression of autophagy-related proteins. Relative expression levels of (A) phospho-Beclin-1, (B) total Beclin-1, (C) p62, (D) LC3A/B in placental tissues from rats exposed to chronic intermittent hypoxia (CIH) or Normoxia during gestation (gestational days 15-20). (E) are representative images of Western blot images for Beclin-1, p62, and LC3A/B. Protein expression levels for phosphorylated forms were calculated as the ratio of phosphorylated levels to total protein, normalized to total loaded protein from Ponceau staining, and expressed as a percentage of Normoxia levels (%control). Total protein levels were normalized similarly. No differences were observed between groups for phospho-Beclin-1 and LC3A/B (p>0.05). Expression levels of total Beclin-1 and p62 were higher in the CIH compared to the Normoxia group. *p=0.006, **p=0.023. Unpaired t-tests were used for all comparisons; Normoxia, n=5-6; CIH, n=5-6), and Welch’s t-tests were used for data with unequal variances (phospho-p38 MAPK; Normoxia, n=6; CIH, n=6). Data are presented as means ± SD. Black square symbols indicate data from the single placenta associated with a female fetus; all other data points represent male fetuses.

#### Decidual responses to CIH

To contextualize the placental stress responses to CIH, we evaluated the decidua, which is a maternal tissue in close anatomical and functional proximity to the placenta. In contrast to placental responses, CIH changed mRNA expression of cytokines in decidual tissues.

Specifically, CIH increased *tnfα* (Mann-Whitney *U* test, p=0.004, Figure 10A), reduced *il-1β* (Unpaired t-test, p=0.053, Figure 10B) and did not affect *il-10* (Mann-Whitney *U* test, p=0.179; Figure 10C). Furthermore, there were no group differences in decidual *il-6* (2^-ΔΔCt^, Normoxia (n=6): 1.1 ± 0.50 vs. CIH (n=6): 1.26 ± 0.90, Mann-Whitney *U* test, p=0.818) and *il-18* (2^-ΔΔCt^, Normoxia (n=6): 1.5 ± 0.23 vs. CIH (n=6): 2.56 ± 1.73, Mann-Whitney *U* test, p=0.662).*<ι><ιτ></i>* There were no differences in phosphorylated and total protein levels of p38 MAPK (Unpaired t-test, p=0.301 and Mann-Whitney *U* test, p=0.815, respectively, n=6/group for all; average data/group not shown; Supplementary Materials, Figure S11) or p44/42 MAPK (Unpaired t-test, p=0.288 and Mann-Whitney *U* test, p=0.485, n=6/group for all; average data/group not shown; Supplementary Materials, Figure S12) of decidual tissues in response to CIH. On the other hand, CIH increased p62 expression (Mann-Whitney *U* test, p=0.026, Figure 10D & Supplementary Materials, Figure S13), did not affect phosphorylation of Beclin-1 (Unpaired t-test, p=0.254, Figure 10E & Supplementary Materials, Figure S14), and increased total Beclin expression (Mann-Whitney *U* test, p=0.015, Figure 10F & Supplementary Materials, Figure S14), while it did not affect LC3A/B (Unpaired t-test, p=0.621, n=6/group; average data/group not shown; Supplementary Materials, Figure S15) in decidual tissues. Representative Western blot images of p62 and Beclin-1 are shown in Figure 10F.

**Figure 10.**
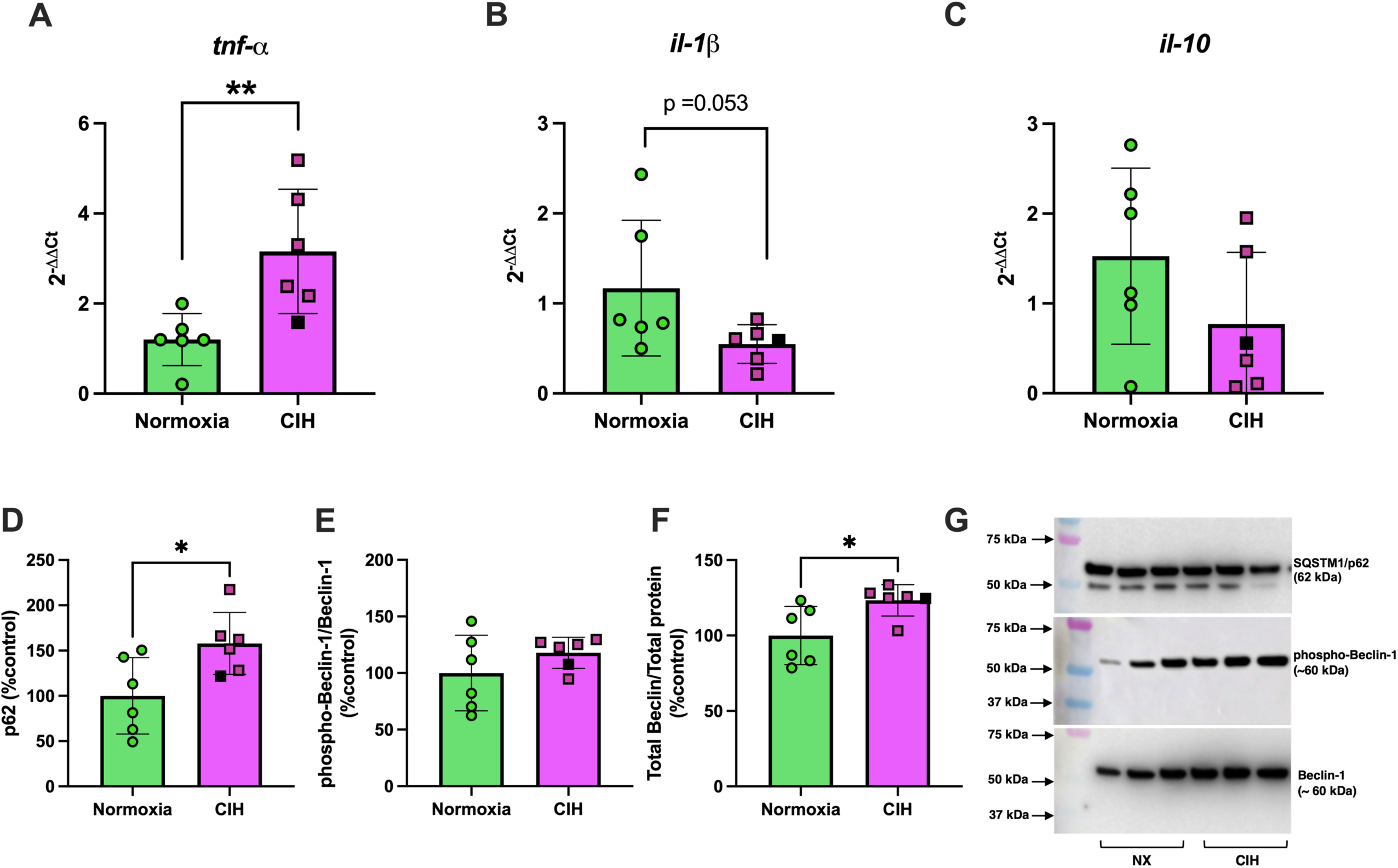
Decidual cytokine mRNA and autophagy marker protein expression. Relative mRNA expression levels (2^-ΔΔCt^) of (A) tumor necrosis factor alpha (*tnf-α*), (B) interleukin-1β (*il-1β*), and relative protein expression levels of (C) p62, (D) phospho-Beclin-1/total Beclin-1, (E) total Beclin-1 in decidual tissues from rats exposed to chronic intermittent hypoxia (CIH) or Normoxia during gestation (gestational days 15-20). (F) are representative images of Western blot images for p62 and Beclin-1. Protein expression levels for phosphorylated forms were calculated as the ratio of phosphorylated levels to total protein, normalized to total loaded protein from Ponceau staining, and expressed as a percentage of Normoxia levels (%control). Total protein levels were normalized similarly. mRNA levels of *tnf-α* (**p=0.004) and total Beclin-1 (p=0.015) were greater, while mRNA levels of *il-1β* were lower (p=0.053) in CIH compared to normoxia groups. Unpaired t-tests were used for normally distributed data with equal variances (*il-1β*, phospho-Beclin-1/Beclin-1; Normoxia, n=6; CIH, n=6), Mann-Whitney *U*-test was used for non-normally distributed data (*tnf-α,* p62, Beclin-1; Normoxia, n=6; CIH, n=6). Data are presented as means ± SD. Black square symbols indicate data from the single decidua associated with a female fetus; all other data points represent male fetuses.

## Discussion

The main findings of this study demonstrate that exposure to CIH during the second half of rat pregnancy 1) induces systemic maternal inflammation, evidenced by an increase in plasma IL-18; 2) reduces concentrations of exosome-like EVs, while leaving EV-mtDNA content unchanged, suggesting a possible increase in mtDNA load per EV; 3) increases placental weights without affecting fetal growth, suggesting reduced fetoplacental efficiency, which is accompanied by upregulation of placental antioxidant enzymes and disruption of autophagy.

Collectively, these findings demonstrate for the first time that even mild, short-term, intermittent hypoxia late in gestation is sufficient to trigger a maternal inflammatory response and to induce both phenotypic and molecular adaptations in the placenta, despite occurring after placental development is largely complete.

### Maternal responses to gestational CIH

CIH increased maternal hematocrit and hemoglobin concentrations, which is a typical response to systemic hypoxia that indicates a reduction in maternal oxygen sensing (25). These hematologic adaptations are driven by a hypoxia-induced increase in erythropoietin levels and an increase in red blood cell production (41). Gestational hypoxia induces changes in maternal caloric intake and body weight gain during rodent pregnancy (41, 42), and the magnitude of these effects depend on hypoxia severity, duration, and onset of the hypoxic stimulus. In the present study, dams increased their food intake on the first day of exposure to CIH, but this change was transient as food intake did not differ between Normoxia and CIH groups during the rest of gestation. CIH did not affect water intake but blunted normal gestational weight gain from GD15 to GD20. The reductions in maternal body weight gain and the increases in maternal hematocrit indicate that a 5-day CIH exposure sufficed to reduce maternal weight gain during pregnancy.

*In vivo* exposure to hypoxic stimuli, including CIH, induces systemic and local inflammatory responses (13, 24, 27). In non-pregnant female rats, CIH exposure for 14 days elicits a systemic proinflammatory response, evidenced by an increase in the IL-6 to IL-10 ratio (27). In pregnant patients with obstructive sleep apnea and gestational diabetes, there is a direct association between intermittent hypoxic episodes and systemic inflammation, quantified as higher concentrations of TNF-α and IL-1β in the maternal circulation (43). Previous studies have implicated IL-18 expression in conditions of systemic and local hypoxia (44–46). In the present study, we observed an increase in the proinflammatory cytokine IL-18 in dams exposed to CIH during late pregnancy. An increase in IL-18 was also observed when primary human trophoblast cells from healthy, late term pregnancies (37-40 weeks) were exposed to chronic hypoxia for 72 hours (44). Additionally, *in vivo* studies comparing rats with preeclampsia-like signs and placental ischemia and healthy controls demonstrated an increase in circulating IL-18 expression (46). Mechanistically, expression of IL-18 is associated with activation of the NLRP3 inflammasome pathway (47), and NLRP3 inflammasome activation has been demonstrated to be increased in pregnancies with preeclampsia (44, 45, 48).

Circulating cell-free mtDNA, released in response to cellular stress and tissue damage, functions as a damage-associated molecular pattern that is recognized by the innate immune system and can trigger inflammatory signaling pathways (32). In non-obstetric conditions associated with oxidative stress and inflammation, such as COVID-19 infections and cardiovascular diseases, ccf-mtDNA has been used as a non-invasive clinical marker to assess disease severity (49, 50) and may serve as a prognostic indicator in hospitalized patients (49, 51). The relevance of ccf-mtDNA in pregnancy complications has been previously reviewed in (32). In addition, we and others have also demonstrated the potential utility of ccf-mtDNA in predictive models for identifying pregnancies at risk of developing adverse outcomes, such as preeclampsia (17, 33). In the present study, we found no increase in content of ccf-mtDNA or systemic oxidative stress in response to CIH during late gestation. These findings may suggest a dependence between these two variables. It is noteworthy that our measurements of ccf-mtDNA reflect all membrane-bound forms of ccf-mtDNA, including those encapsulated in EVs. Therefore, the impact of CIH on the concentration, content, and characteristics of membrane-bound structures carrying mtDNA, including EVs, should be considered when interpreting ccf-mtDNA responses to CIH.

Recently, we demonstrated that mtDNA is released from trophoblast cells in response to oxidative stress in both membrane-bound and non-membrane-bound forms, including exosome-like EVs (16, 17). In obstetric complications, EV characteristics, including size, concentration, and cargo, are altered compared to those from healthy pregnancies (52). While we observed no change in total circulating ccf-mtDNA, we detected a reduction in circulating EV concentrations. EV-mtDNA content, as measured relative to vesicle number, was unchanged, suggesting a potential increase in mtDNA load per EV under CIH conditions. These findings imply that CIH may alter the packaging or release dynamics of mtDNA-containing EVs without changing the overall quantity of circulating membrane-bound mtDNA. This hypothesis is further supported by our previous publications demonstrating that mitochondrial dysregulation in human placenta is associated with increased EV production in pregnancies complicated by hypertensive disorders of pregnancy (53).

Collectively, our data suggest that gestational CIH induces systemic maternal inflammation, manifested by an increase in circulating IL-18 concentrations and mtDNA load in circulating exosome-like EVs. We postulate that activation of the NLRP-3 inflammasome pathway may contribute to CIH-induced maternal inflammation.

### Placental growth and stress responses to CIH

Gestational hypoxia is often associated with fetal growth restriction and adverse fetal outcomes in humans (1), as well as in rodent (9) and ovine models (2). The placenta adapts to changes in nutrients and oxygenation by changing placental structure, size, and function.

Importantly, placental adaptations may reflect adverse effects on fetal development, the magnitude of which depends on timing and severity of insult (9). In the present study, placentas from CIH-exposed dams had increased weight, in the absence of any changes in fetal weight, suggesting a reduction in placental efficiency. Previous studies have shown that placental efficiency, an indicator of fetal development and nutrient exchange, varies in response to changes in environmental conditions, such as hypoxia (54). Given the predominance of male placental samples, this adaptation may represent a male-specific response.

A reduction in placental efficiency, in the absence of fetal weight changes, has been previously reported in studies exposing dams to chronic, sustained hypoxia (13% O_2_) from GD6 to GD20 (41, 55). In contrast, a more severe sustained hypoxic insult (9% O_2_) during late gestation (GD14.5 to GD17.5) reduced both placental and fetal weights (56). The influence of hypoxia severity has been also demonstrated in studies using intermittent rather than sustained hypoxia protocols during pregnancy. In a recent study, Weng et al. reported that moderate CIH (21-10% O_2_, 20 cycles/hour for a total of 8 hours) from GD0 to GD19 had no effect on fetoplacental biometrics, while severe CIH (21-5% O_2_, same cycling pattern) of the same duration resulted in decreased placental and fetal weights (18).

Taken together, these findings suggest that not only the severity and length of hypoxic insults but also their onset relative to placental development are critical (57). Hypoxia beginning early in gestation, while the placenta is still forming, may hinder subsequent growth, whereas hypoxic insults imposed after the placenta is established may impair its functional efficiency or damage previously healthy tissue.

In addition to placental changes in function and size, intermittent fluctuations in oxygen delivery from the maternal circulation into the intervillous space, where maternal-fetal gas exchange occurs, can trigger a placental cell stress response, characterized by redox imbalance and inflammation (13, 58). In pregnant mice, CIH exposure from the beginning of pregnancy until GD14.5, a period that includes early placental development, caused placental oxidative stress and cell death (13). We did not observe cell death in placentas from rats exposed to a shorter CIH protocol (GD15-20), which is consistent with our previous findings (19). Furthermore, we found no change in placental cytokine expression or activation of MAPK signaling proteins, which are known to regulate cellular stress processes, including oxidative stress and inflammation (38, 39). In contrast, cellular defense mechanisms, such as antioxidant enzymes and autophagy, were significantly impacted by CIH.

We observed an increase in placental expression of *catalase* and *sod2* in response to CIH but no change in protein abundance of these enzymes. An increase in mRNA levels in the absence of protein changes during CIH suggest that although transcription of antioxidant enzymes is upregulated, post-transcriptional regulation or protein turnover may limit changes in protein abundance to cope with oxidative stress. Previous studies have shown that a reduction in *sod2* mRNA is associated with reduced activity of SOD2 and increased mitochondrial oxidative stress, indicating a correlation between enzyme gene expression and activity (59).

Others have demonstrated that a 14-day exposure to CIH did not affect SOD1, SOD2, or Catalase in diaphragm muscle in wild-type mice, but it increased mRNA expression of these enzymes in NADPH Oxidase-2 knock-out mice, suggesting that changes in antioxidant enzymes in response to CIH may be NADPH Oxidase-2 (NOX2) dependent (60). Although NOX2 activity was not directly assessed in this study, future studies examining the involvement of NOX2 and mitochondrial reactive oxygen species production in regulating placental antioxidant responses to CIH are warranted.

To further characterize the placental cell stress response, we assessed markers of autophagy, a process that plays a critical role in cellular homeostasis and have been implicated in hypoxia-induced stress adaptations (40). We observed increased protein expression of total Beclin-1 and p62, while phosphorylated Beclin-1 and LC3A/B protein levels remained unchanged. Elevated p62 accumulation, without a corresponding increase in LC3A/B or phospho-Beclin-1, may indicate that gestational CIH in late pregnancy impairs autophagic flux or dysregulates degradation dynamics, warranting further mechanistic investigation (61–64).

The decidua is a specialized, highly vascularized lining of the uterus, where the embryo implants and receives its initial nourishment until a placenta is formed (65). In the present study, we evaluated the decidua as a maternal reference tissue in close anatomic proximity to the fetoplacental interface. The process of placentation and the organization and structure of placental sites, including hemochorial architecture, the presence of distinct functional layers (including the decidua), and deep trophoblast invasion, exhibit significant similarities between humans and rats (66). In response to CIH, we observed an increase in *tnfα* mRNA, a reduction in *il-1β* mRNA and an increase in p62 and Beclin-1 protein expression, while no changes were detected in LC3A/B protein levels.

Notably, the inflammatory response we observed in the decidua, but not in the placenta, may reflect distinct cellular composition and function of these two tissues (67). The decidua contains a substantial population of immune cells, particularly uterine natural killer cells, macrophages, and T cells, which represent the second most abundant cell type after stromal cells. In contrast, the placenta is primarily composed of trophoblast cells specialized for nutrient exchange and immune modulation, which may account for its more restrained inflammatory response to CIH. Furthermore, the inflammatory response of the decidua, which is a maternal tissue, may parallel the systemic inflammatory response we observed in dams exposed to CIH.

Consistent with the response we observed in placental tissues, we noted an increase in p62 and total Beclin-1 expression, in the absence of changes in LC3A/B in deciduae. These data suggest that CIH may induce partial initiation of autophagy; however, confirmation of this possibility would require assessment of dynamic autophagy markers, such as LC3-II accumulation with or without lysosomal inhibitors, in both decidual and placental tissues.

Like placentas, decidual samples tested in this study were predominantly from pregnancies carrying male fetuses, indicating that the observed inflammatory and autophagy responses in decidua may represent male-specific adaptations.

## Strengths and Limitations

We used a rat model of gestational CIH that we previously established to have adverse effects on offspring neural and behavioral outcomes (19, 20). Building on our prior observations that gestational CIH does not affect markers of oxidative stress in the maternal circulation or placenta (19), the current study demonstrates changes in placental antioxidant defense mechanisms, suggesting that the placenta may activate compensatory pathways to maintain redox balance (68). Due to the complexity of the redox system in maintaining cellular homeostasis, further studies are needed to confirm this hypothesis.

A strength of this study is the characterization and quantification of EV-mtDNA in maternal circulation under gestational hypoxic conditions. However, we did not determine the specific source of these EVs. Previous studies from our laboratory have demonstrated mtDNA can be stored within vesicular structures (17) and released from trophoblast cells in response to oxidative stress (16). Future studies using placenta-specific markers, such as placental alkaline phosphatase (69), could assess the direct contribution of placental EVs to the circulating mtDNA pool.

Although fetal sex was determined in this study, nearly all placentas analyzed were from male fetuses (except one), which limits our ability to draw conclusions about sex-specific effects. Thus, the findings of the present study may primarily reflect placental responses to CIH in pregnancies carrying male fetuses. We previously showed that CIH induced sex-dependent alterations in subcortical brain maturation in offspring, with long-term motor and memory deficits observed in male offspring, while females exhibited distinct behavioral impairments (19, 20).

These findings, along with growing evidence of sex differences in placental adaptations to maternal stress (70), highlight the importance of future studies designed with balanced representation of fetal sex.

## Conclusions and Perspectives

In conclusion, our study demonstrates that exposure to gestational CIH during late pregnancy induces maternal inflammation and alters placental antioxidant and autophagy-related responses affecting fetal growth. While total ccf-mtDNA remained unchanged, the reduced concentration of EVs suggests altered packaging or release dynamics of mtDNA in response to CIH. Despite molecular changes in the placenta, we found no differences in fetal growth, litter size, or fetal hematocrit between CIH- and normoxia-exposed groups. Thus, one could speculate that fetal development was preserved under CIH conditions. However, we have recently reported that offspring from pregnancies exposed to CIH exhibit neurodevelopmental and behavioral deficits at various stages of postnatal development (i.e., puberty, young adulthood) (19, 20). Therefore, we posit that CIH impairs fetal organ development and program long-term offspring consequences in the absence of measurable changes in fetal size in late pregnancy. Importantly, our current findings suggest that molecular alterations and size adaptations in the placenta as well as changes in EV concentrations and cargo may serve as early indicators of these long-term consequences.

## Acknowledgements

We acknowledge Johnathan Tune, Ph.D. and Gregory Dick, Ph.D. at the Department of Physiology and Anatomy at the University of North Texas Health Science Center at Fort Worth for equipment use of the blood gas data analysis. Present address of Selina M. Tucker: Fetal Health Center, Children’s Mercy Research Institute, Children’s Mercy Kansas City, Kansas City, MO, USA. Present address of Steve Mabry: North Texas Eye Research Institute, University of North Texas Health Science Center, Fort Worth, TX, USA.

## Author Contributions

SG conceptualized the study. JG, JLB, NRP, RLC, RNOS, and SG designed the study and experiments. DE, ENW, IG, JG, JLB, LL, NH, RLC, RNOS, SM, and ST collected data. IG, JG, JLB, NH, NRP, RLC, RNOS, SG, and ST analyzed data and performed statistical analysis. DE, JG, NH, RNOS, and SG prepared figures. JG, RNOS, and SG drafted manuscript. All authors contributed to data interpretation and edited the manuscript. All authors approved the final version of the manuscript.

## References

1. Colson A, Sonveaux P, Debieve F, and Sferruzzi-Perri AN. Adaptations of the human placenta to hypoxia: opportunities for interventions in fetal growth restriction. Hum Reprod Update 27: 531–569, 2021.

2. Tong W, Allison BJ, Brain KL, Patey OV, Niu Y, Botting KJ, Ford SG, Garrud TA, Wooding PFB, Shaw CJ, Lyu Q, Zhang L, Ma J, Cindrova-Davies T, Yung HW, Burton GJ, and Giussani DA. Chronic Hypoxia in Ovine Pregnancy Recapitulates Physiological and Molecular Markers of Preeclampsia in the Mother, Placenta, and Offspring. Hypertension 79: 1525–1535, 2022.

3. Cresswell JA, Alexander M, Chong MYC, Link HM, Pejchinovska M, Gazeley U, Ahmed SMA, Chou D, Moller AB, Simpson D, Alkema L, Villanueva G, Sguassero Y, Tuncalp O, Long Q, Xiao S, and Say L. Global and regional causes of maternal deaths 2009-20: a WHO systematic analysis. Lancet Glob Health 13: e626–e634, 2025.

4. Larsen LG, Clausen HV, and Jonsson L. Stereologic examination of placentas from mothers who smoke during pregnancy. Am J Obstet Gynecol 186: 531–537, 2002.

5. Rogers JM. Smoking and pregnancy: Epigenetics and developmental origins of the metabolic syndrome. Birth Defects Res 111: 1259–1269, 2019.

6. Clifton VL, Giles WB, Smith R, Bisits AT, Hempenstall PA, Kessell CG, and Gibson PG. Alterations of placental vascular function in asthmatic pregnancies. Am J Respir Crit Care Med 164: 546–553, 2001.

7. D’Souza R, Ashraf R, Rowe H, Zipursky J, Clarfield L, Maxwell C, Arzola C, Lapinsky S, Paquette K, Murthy S, Cheng MP, and Malhame I. Pregnancy and COVID-19: pharmacologic considerations. Ultrasound Obstet Gynecol 57: 195–203, 2021.

8. Maniaci A, La Via L, Pecorino B, Chiofalo B, Scibilia G, Lavalle S, and Scollo P. Obstructive Sleep Apnea in Pregnancy: A Comprehensive Review of Maternal and Fetal Implications. Neurol Int 16: 522–532, 2024.

9. Sferruzzi-Perri AN, Lopez-Tello J, and Salazar-Petres E. Placental adaptations supporting fetal growth during normal and adverse gestational environments. Exp Physiol 108: 371–397, 2023.

10. Burton GJ. Oxygen, the Janus gas; its effects on human placental development and function. J Anat 215: 27–35, 2009.

11. Aplin JD, Myers JE, Timms K, and Westwood M. Tracking placental development in health and disease. Nat Rev Endocrinol 16: 479–494, 2020.

12. Ducsay CA, Goyal R, Pearce WJ, Wilson S, Hu XQ, and Zhang L. Gestational Hypoxia and Developmental Plasticity. Physiol Rev 98: 1241–1334, 2018.

13. Badran M, Abuyassin B, Ayas N, and Laher I. Intermittent hypoxia impairs uterine artery function in pregnant mice. J Physiol 597: 2639–2650, 2019.

14. Vaka VR, McMaster KM, Cunningham MW, Jr., Ibrahim T, Hazlewood R, Usry N, Cornelius DC, Amaral LM, and LaMarca B. Role of Mitochondrial Dysfunction and Reactive Oxygen Species in Mediating Hypertension in the Reduced Uterine Perfusion Pressure Rat Model of Preeclampsia. Hypertension 72: 703–711, 2018.

15. Redman CWG, Staff AC, and Roberts JM. Syncytiotrophoblast stress in preeclampsia: the convergence point for multiple pathways. Am J Obstet Gynecol 226: S907–S927, 2022.

16. Gardner JJ, Cushen SC, Oliveira da Silva RN, Bradshaw JL, Hula N, Gorham IK, Tucker SM, Zhou Z, Cunningham RL, Phillips NR, and Goulopoulou S. Oxidative stress induces release of mitochondrial DNA into the extracellular space in human placental villous trophoblast BeWo cells. Am J Physiol Cell Physiol 2024.

17. Cushen SC, Ricci CA, Bradshaw JL, Silzer T, Blessing A, Sun J, Zhou Z, Scroggins SM, Santillan MK, Santillan DA, Phillips NR, and Goulopoulou S. Reduced Maternal Circulating Cell-Free Mitochondrial DNA Is Associated With the Development of Preeclampsia. J Am Heart Assoc 11: e021726, 2022.

18. Weng C, Huang L, Feng H, He Q, Lin X, Jiang T, Lin J, Wang X, and Liu Q. Gestational chronic intermittent hypoxia induces hypertension, proteinuria, and fetal growth restriction in mice. Sleep Breath 26: 1661–1669, 2022.

19. Wilson EN, Mabry S, Bradshaw JL, Gardner JJ, Rybalchenko N, Engelland R, Fadeyibi O, Osikoya O, Cushen SC, Goulopoulou S, and Cunningham RL. Gestational hypoxia in late pregnancy differentially programs subcortical brain maturation in male and female rat offspring. Biol Sex Differ 13: 54, 2022.

20. Mabry S, Wilson EN, Bradshaw JL, Gardner JJ, Fadeyibi O, Vera E, Jr., Osikoya O, Cushen SC, Karamichos D, Goulopoulou S, and Cunningham RL. Sex and age differences in social and cognitive function in offspring exposed to late gestational hypoxia. Biol Sex Differ 14: 81, 2023.

21. Wilson EN, Anderson M, Snyder B, Duong P, Trieu J, Schreihofer DA, and Cunningham RL. Chronic intermittent hypoxia induces hormonal and male sexual behavioral changes: Hypoxia as an advancer of aging. Physiol Behav 189: 64–73, 2018.

22. Snyder B, Duong P, Tenkorang M, Wilson EN, and Cunningham RL. Rat Strain and Housing Conditions Alter Oxidative Stress and Hormone Responses to Chronic Intermittent Hypoxia. Front Physiol 9: 1554, 2018.

23. Snyder B, Duong P, Trieu J, and Cunningham RL. Androgens modulate chronic intermittent hypoxia effects on brain and behavior. Horm Behav 106: 62–73, 2018.

24. Snyder B, Shell B, Cunningham JT, and Cunningham RL. Chronic intermittent hypoxia induces oxidative stress and inflammation in brain regions associated with early-stage neurodegeneration. Physiol Rep 5: 2017.

25. McGuire M, and Bradford A. Chronic intermittent hypoxia increases haematocrit and causes right ventricular hypertrophy in the rat. Respir Physiol 117: 53–58, 1999.

26. Bradshaw JL, Wilson EN, Gardner JJ, Mabry S, Tucker SM, Rybalchenko N, Vera E, Jr., Goulopoulou S, and Cunningham RL. Pregnancy-induced oxidative stress and inflammation are not associated with impaired maternal neuronal activity or memory function. Am J Physiol Regul Integr Comp Physiol 2024.

27. Mabry S, Bradshaw JL, Gardner JJ, Wilson EN, and Cunningham RL. Sex-dependent effects of chronic intermittent hypoxia: implication for obstructive sleep apnea. Biol Sex Differ 15: 38, 2024.

28. Nicklas JA, Brooks EM, Hunter TC, Single R, and Branda RF. Development of a quantitative PCR (TaqMan) assay for relative mitochondrial DNA copy number and the common mitochondrial DNA deletion in the rat. Environ Mol Mutagen 44: 313–320, 2004.

29. Dhakal P, and Soares MJ. Single-step PCR-based genetic sex determination of rat tissues and cells. Biotechniques 62: 232–233, 2017.

30. Osikoya O, Ahmed H, Panahi S, Bourque SL, and Goulopoulou S. Uterine perivascular adipose tissue is a novel mediator of uterine artery blood flow and reactivity in rat pregnancy. J Physiol 597: 3833–3852, 2019.

31. Bradshaw JL, Cushen SC, Ricci CA, Tucker SM, Gardner JJ, Little JT, Osikoya O, and Goulopoulou S. Exposure to unmethylated CpG oligonucleotides disrupts blood pressure circadian rhythms and placental clock gene network in pregnant rats. Am J Physiol Heart Circ Physiol 325: H323–H337, 2023.

32. Bradshaw JL, Cushen SC, Phillips NR, and Goulopoulou S. Circulating Cell-Free Mitochondrial DNA in Pregnancy. Physiology (Bethesda) 37: 0, 2022.

33. Busnelli A, Lattuada D, Ferrari S, Reschini M, Colciaghi B, Somigliana E, Fedele L, and Ferrazzi E. Mitochondrial DNA Copy Number in Peripheral Blood in the First Trimester of Pregnancy and Different Preeclampsia Clinical Phenotypes Development: A Pilot Study. Reprod Sci 26: 1054–1061, 2019.

34. De Gaetano A, Solodka K, Zanini G, Selleri V, Mattioli AV, Nasi M, and Pinti M. Molecular Mechanisms of mtDNA-Mediated Inflammation. Cells 10: 2021.

35. Marschalek J, Wohlrab P, Ott J, Wojta J, Speidl W, Klein KU, Kiss H, Pateisky P, Zeisler H, and Kuessel L. Maternal serum mitochondrial DNA (mtDNA) levels are elevated in preeclampsia - A matched case-control study. Pregnancy Hypertens 14: 195–199, 2018.

36. Qiu C, Hevner K, Enquobahrie DA, and Williams MA. A case-control study of maternal blood mitochondrial DNA copy number and preeclampsia risk. Int J Mol Epidemiol Genet 3: 237–244, 2012.

37. West AP, and Shadel GS. Mitochondrial DNA in innate immune responses and inflammatory pathology. Nat Rev Immunol 17: 363–375, 2017.

38. Cargnello M, and Roux PP. Activation and function of the MAPKs and their substrates, the MAPK-activated protein kinases. Microbiol Mol Biol Rev 75: 50–83, 2011.

39. Koinzer S, Reinecke K, Herdegen T, Roider J, and Klettner A. Oxidative Stress Induces Biphasic ERK1/2 Activation in the RPE with Distinct Effects on Cell Survival at Early and Late Activation. Curr Eye Res 40: 853–857, 2015.

40. Bellot G, Garcia-Medina R, Gounon P, Chiche J, Roux D, Pouyssegur J, and Mazure NM. Hypoxia-induced autophagy is mediated through hypoxia-inducible factor induction of BNIP3 and BNIP3L via their BH3 domains. Mol Cell Biol 29: 2570–2581, 2009.

41. Nuzzo AM, Camm EJ, Sferruzzi-Perri AN, Ashmore TJ, Yung HW, Cindrova-Davies T, Spiroski AM, Sutherland MR, Logan A, Austin-Williams S, Burton GJ, Rolfo A, Todros T, Murphy MP, and Giussani DA. Placental Adaptation to Early-Onset Hypoxic Pregnancy and Mitochondria-Targeted Antioxidant Therapy in a Rodent Model. Am J Pathol 188: 2704–2716, 2018.

42. Iqbal W, and Ciriello J. Effect of maternal chronic intermittent hypoxia during gestation on offspring growth in the rat. Am J Obstet Gynecol 209: 564 e561–569, 2013.

43. Serednytskyy O, Alonso-Fernandez A, Ribot C, Herranz A, Alvarez A, Sanchez A, Rodriguez P, Gil AV, Pia C, Cubero JP, Barcelo M, Cerda M, Codina M, M DP, Barcelo A, Iglesias A, Morell-Garcia D, Pena JA, Gimenez MP, Pinas MC, and Garcia-Rio F. Systemic inflammation and sympathetic activation in gestational diabetes mellitus with obstructive sleep apnea. BMC Pulm Med 22: 94, 2022.

44. Cheng SB, Nakashima A, Huber WJ, Davis S, Banerjee S, Huang Z, Saito S, Sadovsky Y, and Sharma S. Pyroptosis is a critical inflammatory pathway in the placenta from early onset preeclampsia and in human trophoblasts exposed to hypoxia and endoplasmic reticulum stressors. Cell Death Dis 10: 927, 2019.

45. Nunes PR, Mattioli SV, and Sandrim VC. NLRP3 Activation and Its Relationship to Endothelial Dysfunction and Oxidative Stress: Implications for Preeclampsia and Pharmacological Interventions. Cells 10: 2021.

46. Taylor EB, George EM, Ryan MJ, Garrett MR, and Sasser JM. Immunological comparison of pregnant Dahl salt-sensitive and Sprague-Dawley rats commonly used to model characteristics of preeclampsia. Am J Physiol Regul Integr Comp Physiol 321: R125–R138, 2021.

47. Ihim SA, Abubakar SD, Zian Z, Sasaki T, Saffarioun M, Maleknia S, and Azizi G. Interleukin-18 cytokine in immunity, inflammation, and autoimmunity: Biological role in induction, regulation, and treatment. Front Immunol 13: 919973, 2022.

48. Zeng H, Han X, Zhu Z, Yu S, Mei S, Cheng X, Zhang W, Zhang G, and Fang D. Increased uterine NLRP3 inflammasome and leucocyte infiltration in a rat model of preeclampsia. Am J Reprod Immunol 86: e13493, 2021.

49. Hepokoski ML, Odish M, Lam MT, Coufal NG, Rolfsen ML, Shadel GS, Moyzis AG, Sainz AG, Takiar PG, Patel S, Leonard AJ, Samandari N, Hansen E, Trescott S, Nguyen C, Jepsen K, Cutter G, Gillespie MN, Spragg RG, Sasik R, and Ix JH. Absolute quantification of plasma mitochondrial DNA by droplet digital PCR marks COVID-19 severity over time during intensive care unit admissions. Am J Physiol Lung Cell Mol Physiol 323: L84–L92, 2022.

50. Mahmoodpoor A, Mohammadzadeh M, Asghari R, Tagizadeh M, Iranpour A, Rezayi M, Pahnvar AJ, Emamalizadeh B, Sohrabifar N, and Kazeminasab S. Prognostic potential of circulating cell free mitochondrial DNA levels in COVID-19 patients. Mol Biol Rep 50: 10249–10255, 2023.

51. Wiersma M, van Marion DMS, Bouman EJ, Li J, Zhang D, Ramos KS, Lanters EAH, de Groot NMS, and Brundel B. Cell-Free Circulating Mitochondrial DNA: A Potential Blood-Based Marker for Atrial Fibrillation. Cells 9: 2020.

52. Paul N, Sultana Z, Fisher JJ, Maiti K, and Smith R. Extracellular vesicles-crucial players in human pregnancy. Placenta 140: 30–38, 2023.

53. Ricci CA, Reid DM, Sun J, Santillan DA, Santillan MK, Phillips NR, and Goulopoulou S. Maternal and fetal mitochondrial gene dysregulation in hypertensive disorders of pregnancy. Physiol Genomics 55: 275–285, 2023.

54. Fowden AL, Sferruzzi-Perri AN, Coan PM, Constancia M, and Burton GJ. Placental efficiency and adaptation: endocrine regulation. J Physiol 587: 3459–3472, 2009.

55. Richter HG, Camm EJ, Modi BN, Naeem F, Cross CM, Cindrova-Davies T, Spasic-Boskovic O, Dunster C, Mudway IS, Kelly FJ, Burton GJ, Poston L, and Giussani DA. Ascorbate prevents placental oxidative stress and enhances birth weight in hypoxic pregnancy in rats. J Physiol 590: 1377–1387, 2012.

56. Kimball R, Wayment M, Merrill D, Wahlquist T, Reynolds PR, and Arroyo JA. Hypoxia reduces placental mTOR activation in a hypoxia-induced model of intrauterine growth restriction (IUGR). Physiol Rep 3: 2015.

57. Valverde-Perez E, Olea E, Rocher A, Aaronson PI, and Prieto-Lloret J. Effects of gestational intermittent hypoxia on the respiratory system: A tale of the placenta, fetus, and developing offspring. J Sleep Res e14435, 2024.

58. Hung TH, Skepper JN, and Burton GJ. In vitro ischemia-reperfusion injury in term human placenta as a model for oxidative stress in pathological pregnancies. Am J Pathol 159: 1031–1043, 2001.

59. Peng YJ, Nanduri J, Khan SA, Yuan G, Wang N, Kinsman B, Vaddi DR, Kumar GK, Garcia JA, Semenza GL, and Prabhakar NR. Hypoxia-inducible factor 2alpha (HIF-2alpha) heterozygous-null mice exhibit exaggerated carotid body sensitivity to hypoxia, breathing instability, and hypertension. Proc Natl Acad Sci U S A 108: 3065–3070, 2011.

60. Drummond SE, Burns DP, El Maghrani S, Ziegler O, Healy V, and O’Halloran KD. Chronic Intermittent Hypoxia-Induced Diaphragm Muscle Weakness Is NADPH Oxidase-2 Dependent. Cells 12: 2023.

61. Kang R, Zeh HJ, Lotze MT, and Tang D. The Beclin 1 network regulates autophagy and apoptosis. Cell Death Differ 18: 571–580, 2011.

62. Sun Y, Yao X, Zhang QJ, Zhu M, Liu ZP, Ci B, Xie Y, Carlson D, Rothermel BA, Sun Y, Levine B, Hill JA, Wolf SE, Minei JP, and Zang QS. Beclin-1-Dependent Autophagy Protects the Heart During Sepsis. Circulation 138: 2247–2262, 2018.

63. Au AK, Aneja RK, Bayir H, Bell MJ, Janesko-Feldman K, Kochanek PM, and Clark RSB. Autophagy Biomarkers Beclin 1 and p62 are Increased in Cerebrospinal Fluid after Traumatic Brain Injury. Neurocrit Care 26: 348–355, 2017.

64. Loos B, du Toit A, and Hofmeyr JH. Defining and measuring autophagosome flux-concept and reality. Autophagy 10: 2087–2096, 2014.

65. Mori M, Bogdan A, Balassa T, Csabai T, and Szekeres-Bartho J. The decidua-the maternal bed embracing the embryo-maintains the pregnancy. Semin Immunopathol 38: 635–649, 2016.

66. Soares MJ, Chakraborty D, Karim Rumi MA, Konno T, and Renaud SJ. Rat placentation: an experimental model for investigating the hemochorial maternal-fetal interface. Placenta 33: 233–243, 2012.

67. Yang L, Semmes EC, Ovies C, Megli C, Permar S, Gilner JB, and Coyne CB. Innate immune signaling in trophoblast and decidua organoids defines differential antiviral defenses at the maternal-fetal interface. Elife 11: 2022.

68. Burton GJ, and Jauniaux E. Oxidative stress. Best Pract Res Clin Obstet Gynaecol 25: 287–299, 2011.

69. Dragovic RA, Collett GP, Hole P, Ferguson DJ, Redman CW, Sargent IL, and Tannetta DS. Isolation of syncytiotrophoblast microvesicles and exosomes and their characterisation by multicolour flow cytometry and fluorescence Nanoparticle Tracking Analysis. Methods 87: 64–74, 2015.

70. Kalisch-Smith JI, Simmons DG, Dickinson H, and Moritz KM. Review: Sexual dimorphism in the formation, function and adaptation of the placenta. Placenta 54: 10–16, 2017.

