## Supplementary Tables and Figures for "Gestational Chronic Intermittent Hypoxia Triggers Maternal Inflammation and Disrupts Placental Stress Responses"

*Equal contribution

**Table S1. Chemicals and reagents.**

| Chemical / Reagent Name | Catalog # | Vendor, City, State, Country |
|---|---|---|
| β-mercapoethanol | M3148-25 ML | Sigma-Aldrich, Saint Louis, MO, USA |
| Bovine Serum Albumin | A6003-25G | Sigma-Aldrich, Saint Louis, MO, USA |
| DNeasy Blood & Tissue Kit | 69504 | Qiagen, Redwood City, CA, USA |
| dNTP Mix, 10mM | R0192 | Thermo Fisher Scientific, Waltham, MA, USA |
| DreamTaq™ Green DNA polymerase | EP0711 | Thermo Fisher Scientific, Waltham, MA, USA |
| Exo-Check Exosome antibody array | EXORAY200B-4 | System Biosciences, Palo Alto, CA, USA |
| ExoQuick Exosome Precipitation Solution | EXOQ5A-1 | System Biosciences, Palo Alto, CA, USA |
| Extract-N-AMP-Tissue PCR Kit | XNAT2-1KT | Sigma-Aldrich, Saint Louis, MO, USA |
| GreenRuler 100 bp plus DNA ladder ready to use | SM0323 | Thermo Fisher Scientific, Waltham, MA, USA |
| iQ SYBR Green Supermix, 500 x 50 $\mu$l reactions, 12.5 ml (10 x 1.25 ml) | 1708882 | Bio-Rad, Hercules, CA, USA |
| Millipex Rat Cytokine/Chemokine Magnetic Bead Panel | RECYTMAG-65K | MilliporeSigma, Burlington, MA, USA |
| miRNeasy Mini Kit | 217004 | Qiagen, Redwood City, CA, USA |
| NaF (Sodium fluoride, 100 mM) | 201154 | Sigma-Aldrich, Saint Louis, MO, USA |
| $Na_3VO_4$ (Sodium orthovanadate, 100 mM) | S6508 | Sigma-Aldrich, Saint Louis, MO, USA |
| $Na_2H_2P_2O_7$ (Sodium pyrophosphate dibasic, 10 mM) | 71501 | Sigma-Aldrich, Saint Louis, MO, USA |
| Nitrocellulose membrane, 0.45 $\mu$m 30 cm x 3.5 m | 162-0115 | Bio-Rad, Hercules, CA, USA |
| Non-Fat Milk (Blotting-Grade Blocker) | 1706404 | Bio-Rad, Hercules, CA, USA |
| Oligo-dT primers, 100 $\mu$l (0.4 $\mu$g/$\mu$l) | 79237 | Qiagen, Redwood City, CA, USA |
| OxiSelect Advanced Oxidative Protein Products (AOPP) Assay Kit | STA-318 | Cell Biolabs Inc, San Diego, CA, USA |
| PMSF (Phenylmethylsulfonyl fluoride, 100 mM) | 36978 | Thermo Fisher Scientific, Waltham, MA, USA |

| | | |
|---|---|---|
| Protease Inhibitor Cocktail Tablets | P8340-1ML | Sigma-Aldrich, Saint Louis, MO, USA |
| QIAzol Lysis Reagent | 79306 | Qiagen, Redwood City, CA, USA |
| RiboGuard RNase inhibitor, 50,000 U | RG90910K (Part E0126-40D7) | LGC Biosearch Technologies, Middleton, WI, USA |
| Sensiscript RT Kit Reagents | 205213 | Qiagen, Redwood City, CA, USA |
| TaqMan Universal Master Mix II, no UNG | 4440040 | Applied Biosystems, Waltham, MA, USA |
| T-PER™ Tissue Protein Extraction Reagent | 78510 | Thermo Fisher Scientific, Waltham, MA, USA |

**Table S2. Primers for Kdm5c (X chromosome) and Kdm5d (Y chromosome) genes used in rat placenta sex determination.**

| sex | primer | forward | reverse |
|---|---|---|---|
| *X-chromosome* | Kdm5c (F1) | 3'- CACCCCGGATCAGCATGTTT -5' | |
| *Y-chromosome* | Kdm5d (F2) | 3'-CCTTAGTCGGTAGAGTGGTT-5' | |
| *XY chromosome* | Kdm5c + kdm5d | | 5'- CCGCTGCCAAATTCTTTGG -3' |

As detailed in Dhakal et al. (1), we used the following gene sequences for primers of Kdm5c (X chromosome, NC_005120.4) and Kdm5d (Y chromosome, NC_024475.1) for placental sex determination in rat.

**Table S3.** List of forward and reverse primers sequence (5’ to 3’) of target genes for quantitative real time-PCR.

| GENE | FORWARD | REVERSE |
|---|---|---|
| *sod 1* | *TTGGCCGTACTATGGTGGTC* | *GGGCAATCCCAATCACACCA* |
| *sod 2* | *CGGGGGCCATATCAATCACA* | *GCCTCCAGCAACTCTCCTTT* |
| *catalase* | *CTGACTGACGCGATTGCCTA* | *ATGGTGTAGGATTGCGGAGC* |
| *il-6* | *TGATGGATGCTTCCAAACTG* | *GAGCTTGGAAGTTGGGGTA* |
| *il-10* | *CTG GCT CAG CAC TGC TAT GT* | *GTCGTTGCTTGTCTCTCCTTGTA* |
| *il-1*$\beta$ | *ACTGAACTTCGGGGTGATTG* | *GCTTGGTGGTTTGCTACGAC* |
| *tnf-*$\alpha$ | *AGCTCTTCTACCAGCAAACATC* | *GCTTGGTGGTTTGCTACGAC* |
| *il-18* | *AGCTCTTCTACCAGCAAACATC* | *CTTCCAACTGAGAGGCTGTGC* |
| *gapdh* | *AGACAGCCGCATCTTCTTGT* | *TACGGCCAAATCCGTTCACA* |

**Table S4.** Antibody information – Western blots

| Protein | Cat# | Vendor/City, State, Country | Molecular Weight (kDa) | Dilution (primary antibody) | Secondary antibody (dilution; company, cat #) | Blocking buffer | RRIDs |
|---|---|---|---|---|---|---|---|
| Beclin-1 (D40C5) | 3495S | Cell Signaling/Danvers, MA, USA | 60 | 1:1000 | 1:5000; Anti-Rabbit IgG, HRP-Linked Antibody #7074, Cell Signaling | Milk 5% | RRID: AB_1903911 |
| Caspase-3 Control Cell Extracts | 9663S | Cell Signaling/Danvers, MA, USA | 30 | 1:1000 | 1:5000; Anti-Rabbit IgG, HRP-Linked Antibody #7074, Cell Signaling | Milk 5% | N/A |
| Caspase-3 (D3R6Y) | 14220T | Cell Signaling/Danvers, MA, USA | 30 | 1:1000 | 1:5000; Anti-Rabbit IgG, HRP-Linked Antibody #7074, Cell Signaling | Milk 5% | RRID: AB_2798429 |
| Catalase (D5N7V) | 14097S | Cell Signaling/Danvers, MA, USA | 60 | 1:1000 | 1:5000; Anti-Rabbit IgG, HRP-Linked Antibody #7074, Cell Signaling | BSA 3% | RRID: AB_2798391 |
| Cleaved Caspase-3 (Asp175) | 9661S | Cell Signaling/Danvers, MA, USA | 17/19 | 1:500 | 1:2000; Anti-Rabbit IgG, HRP-Linked Antibody #7074, Cell Signaling | Milk 5% | RRID: AB_2341188 |
| LC3A/B | 4108S | Cell Signaling/Danvers, MA, USA | 14/116 | 1:1000 | 1:2000; Anti-Rabbit IgG, HRP-Linked Antibody #7074, Cell Signaling | Milk 5% | RRID: AB_2137703 |
| LC3 Control Cell Extracts (HeLa untreated) | 73774 | Cell Signaling/Danvers, MA, USA | 14/16 | 1:1000 | 1:5000; Anti-Rabbit IgG, HRP-Linked Antibody #7074, Cell Signaling | Milk 5% | N/A |

| | | | | | | | |
|---|---|---|---|---|---|---|---|
| LC3 Control Cell Extracts _ (HeLa +Chloroquine) | 96900 | Cell Signaling/Danvers, MA, USA | 14/16 | 1:1000 | 1:5000; Anti-Rabbit IgG, HRP-Linked Antibody #7074, Cell Signaling | Milk 5% | N/A |
| p38 MAPK | 9212S | Cell Signaling/Danvers, MA, USA | 38 | 1:1000 | 1:2000; Anti-Rabbit IgG, HRP-Linked Antibody #7074, Cell Signaling | Milk 5% | RRID: AB_330713 |
| p38 MAPK Control Cell Extracts (C-6 +Anisomycin) | 48080 S | Cell Signaling/Danvers, MA, USA | 38 | 1:1000 | 1:2000 Anti-Rabbit IgG, HRP-Linked Antibody #7074, Cell Signaling | Milk 5% | N/A |
| p38 MAPK Control Cell Extracts (C-6 untreated) | 28670 S | Cell Signaling/Danvers, MA, USA | 38 | 1:1000 | 1:2000 Anti-Rabbit IgG, HRP-Linked Antibody #7074, Cell Signaling | Milk 5% | N/A |
| p44/42 MAPK (Erk1/2) | 9102S | Cell Signaling/Danvers, MA, USA | 44/42 | 1:1000 | 1:5000; Anti-Rabbit IgG, HRP-Linked Antibody #7074, Cell Signaling | BSA 3% | RRID: AB_330744 |
| Phospho-Beclin-1 (Ser30) (E1C4X) | 35955 S | Cell Signaling/Danvers, MA, USA | 60 | 1:1000 | 1:5000; Anti-Rabbit IgG, HRP-Linked Antibody #7074, Cell Signaling | Milk 5% | N/A |
| Phospho-p44/42 MAPK (Erk1/2) (Thr202/Tyr204) | 9101S | Cell Signaling/Danvers, MA, USA | 44/42 | 1:1000 | 1:5000; Anti-Rabbit IgG, HRP-Linked Antibody #7074, Cell Signaling | BSA 3% | RRID: AB_331646 |
| SOD1 (E4G1H) XP® | 37385 S | Cell Signaling/Danvers, MA, USA | 18 | 1:1000 | 1:5000; Anti-Rabbit IgG, HRP-Linked Antibody #7074, Cell Signaling | BSA 3% | RRID: AB_3073954 |
| SOD2 (D9V9C) | 13194 S | Cell Signaling/Danvers, MA, USA | 22 | 1:1000 | 1:5000; Anti-Rabbit IgG, HRP-Linked Antibody #7074, Cell Signaling | BSA 3% | RRID: AB_2750869 |

| | | | | | | | |
|---|---|---|---|---|---|---|---|
| SQSTM1/p62 | 5114S | Cell Signaling/Danvers, MA, USA | 62 | 1:1000 | 1:5000; Anti-Rabbit IgG, HRP-Linked Antibody #7074, Cell Signaling | Milk 5% | RRID: AB_10624872 |

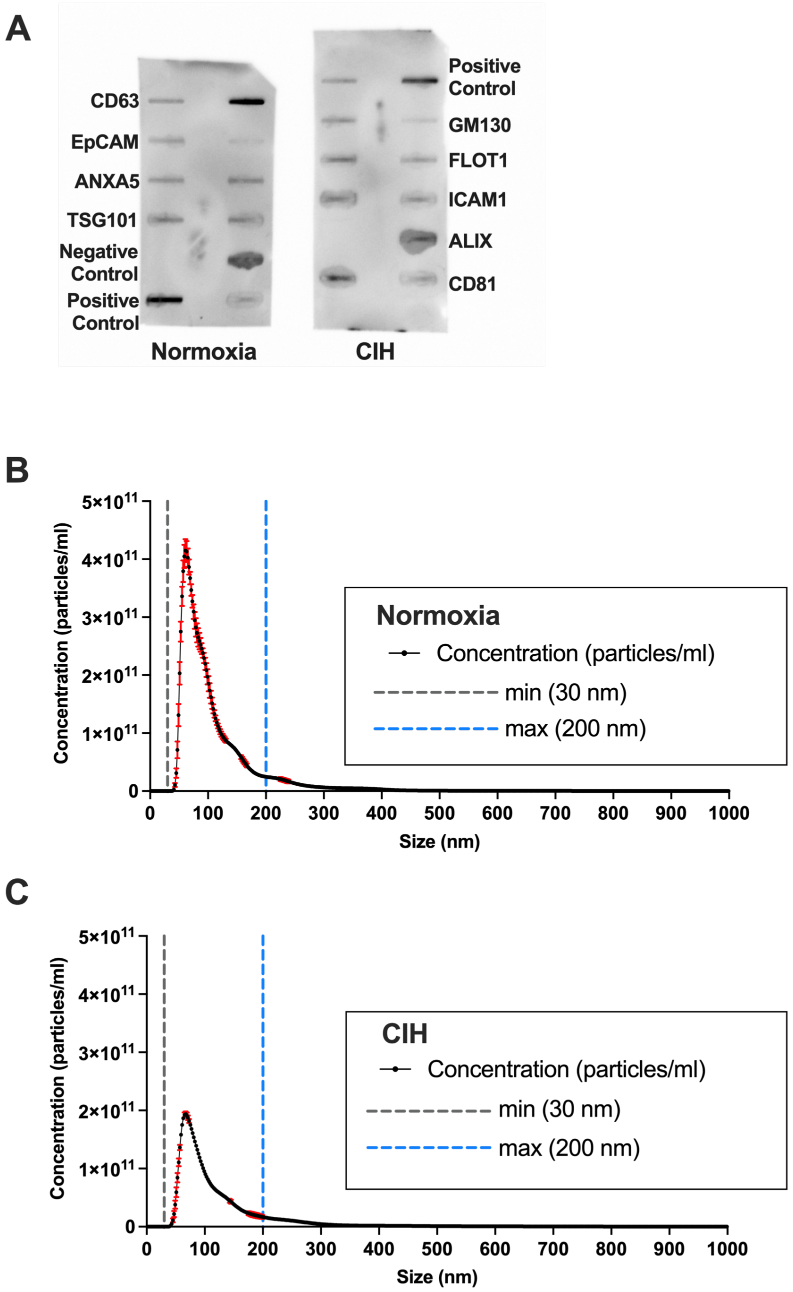


**Figure S1. A)** Exo-check™ representative images. Semi-quantitative Exo-Check™ Exosome Antibody Array was exposed to 50 µg of exosomal proteins isolated from rat serum at gestational day 20 using ExoQuick™. The following exosome markers were used in the array: CD63 and CD81 (tetraspanins), EpCam (epithelial cell adhesion molecule; often found in cancer-derived exosomes), ANX5 (annexin 5), TSG01 (tumor susceptibility gene 101), FLOT-1 (flotillin-1), ICAM-1 (intercellular adhesion molecule 1), ALIX (programmed cell death 6 interacting protein (PDCD6IP). A labeled positive control (labeled HRP detection) and a negative control (blank spot as a background control) have been included. **B-C)** Nanoparticle tracking analysis was used to determine size distribution (nm) of serum EVs from Normoxia and CIH-exposed dams. Each graph depicts the average of

3 readings for each group. The threshold lines at 30 nm (grey dotted line) and 200 nm (blue dotted line) denote the size range of small EVs. Red error bars indicate ± SEM.

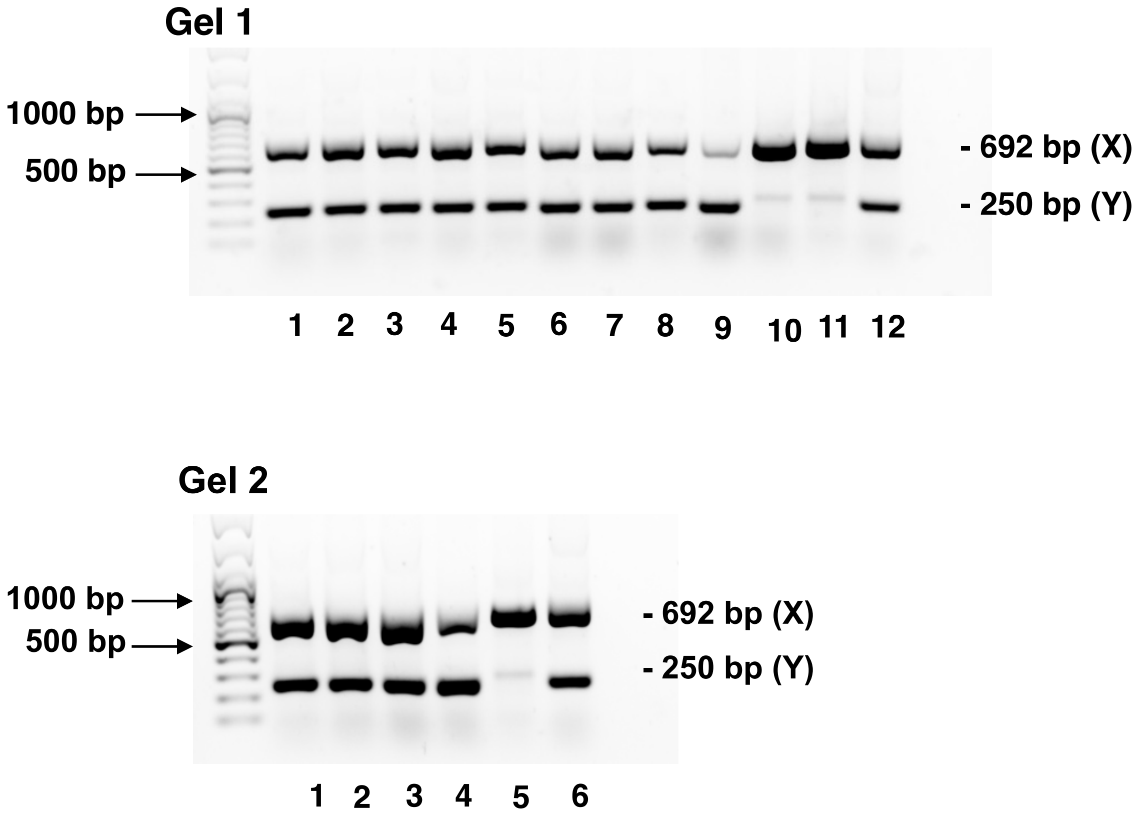


**Figure S2. Fetal placenta PCR products corresponding to X and Y chromosome-specific genes**. As determined in Dhakal et al.(1), we identified males by a band at 250 and 692 bp, and females by a band at 692 bp. Bands were viewed on 1% agarose gel using GeneRuler 100-bp Plus DNA Ladder. Tails from adult male and female rats were used as controls. **Gel 1:** L) GeneRuler 100-bp Plus DNA Ladder; Lanes 1-5: Normoxic rat placenta; Lanes 6-10: CIH rat placenta; Lane 11: Female adult rat tail, control; Lane 12: Male adult rat tail, control. **Gel 2:** L) GeneRuler 100-bp Plus DNA Ladder; Lanes 1-2: Normoxic rat placenta; Lanes 3-4: CIH rat placenta; Lane 5: Female adult rat tail, control; Lane 6: Male adult rat tail, control.

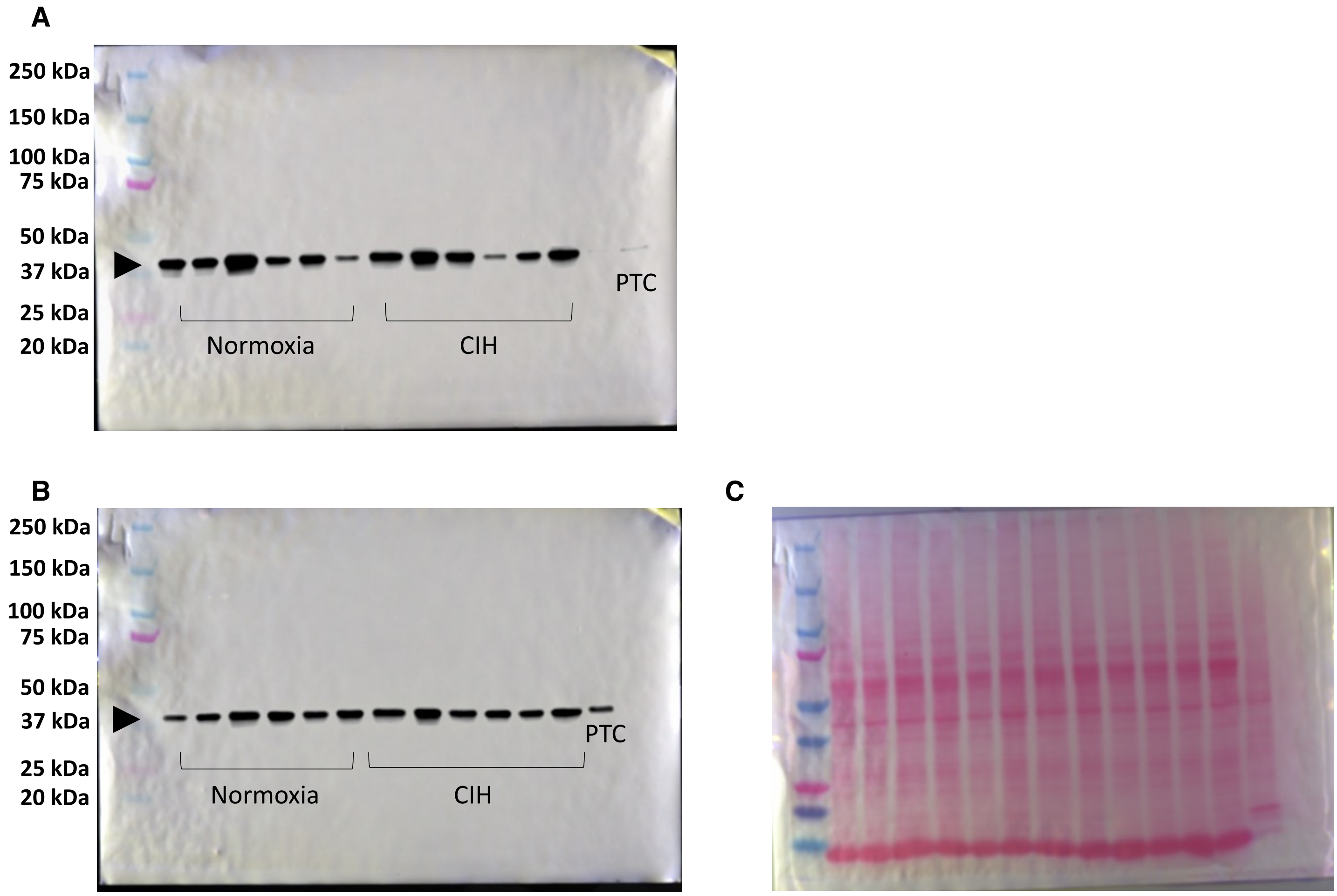


**Figure S3.** Representative immunoblot staining against phospho-p38 (A), p38 (B) and Ponceau S total protein staining (C) in rat Normoxia and chronic intermittent hypoxia (CIH) placentas. C-6 glioma cells treated with anisomycin was used as a positive control (PTC).

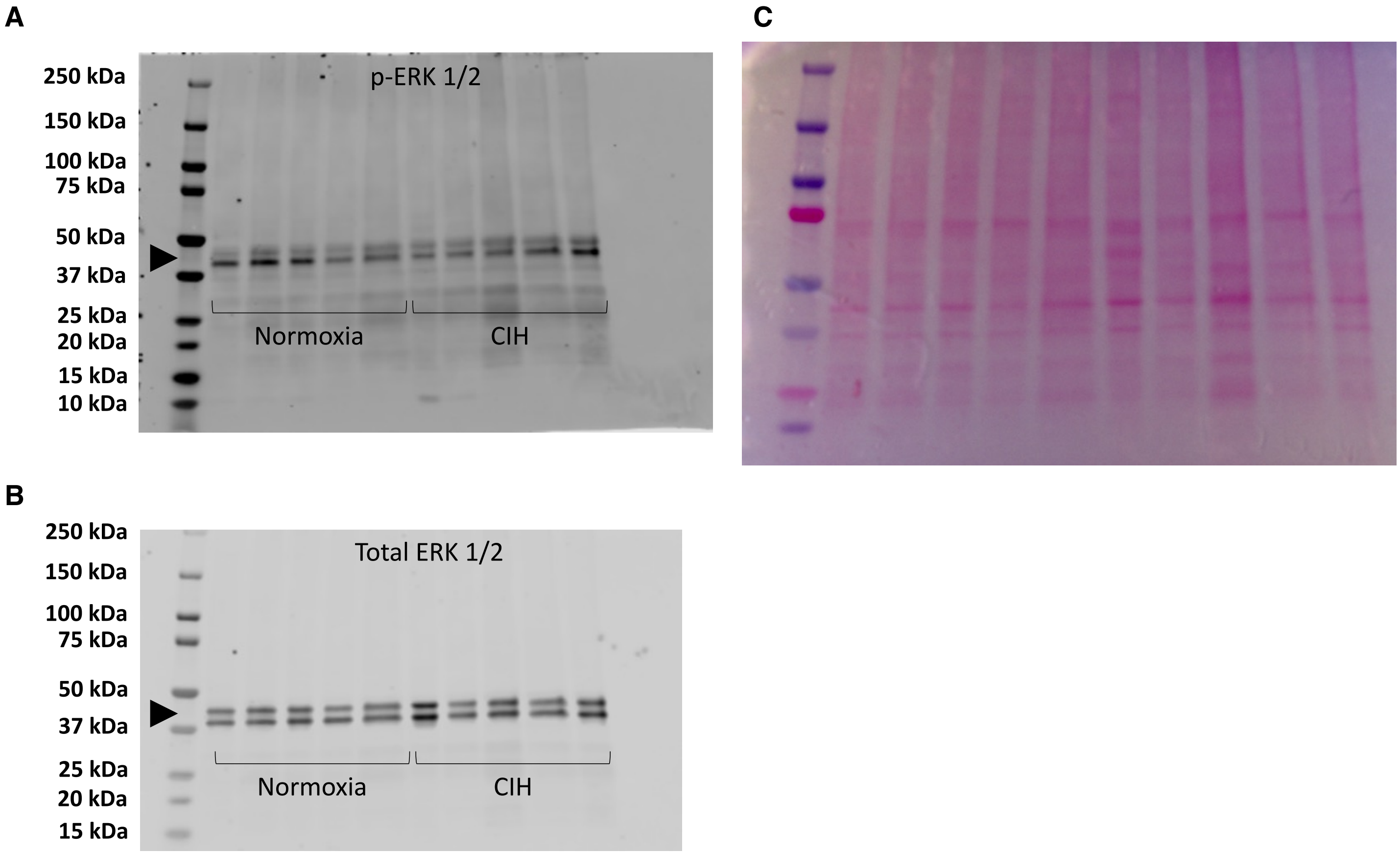


**Figure S4.** Representative immunoblot staining against phosphor-ERK 1/2 (A), ERK 1/2 (B) and Ponceau S total protein staining (C) in rat normoxia and chronic intermittent hypoxia (CIH) placentas.

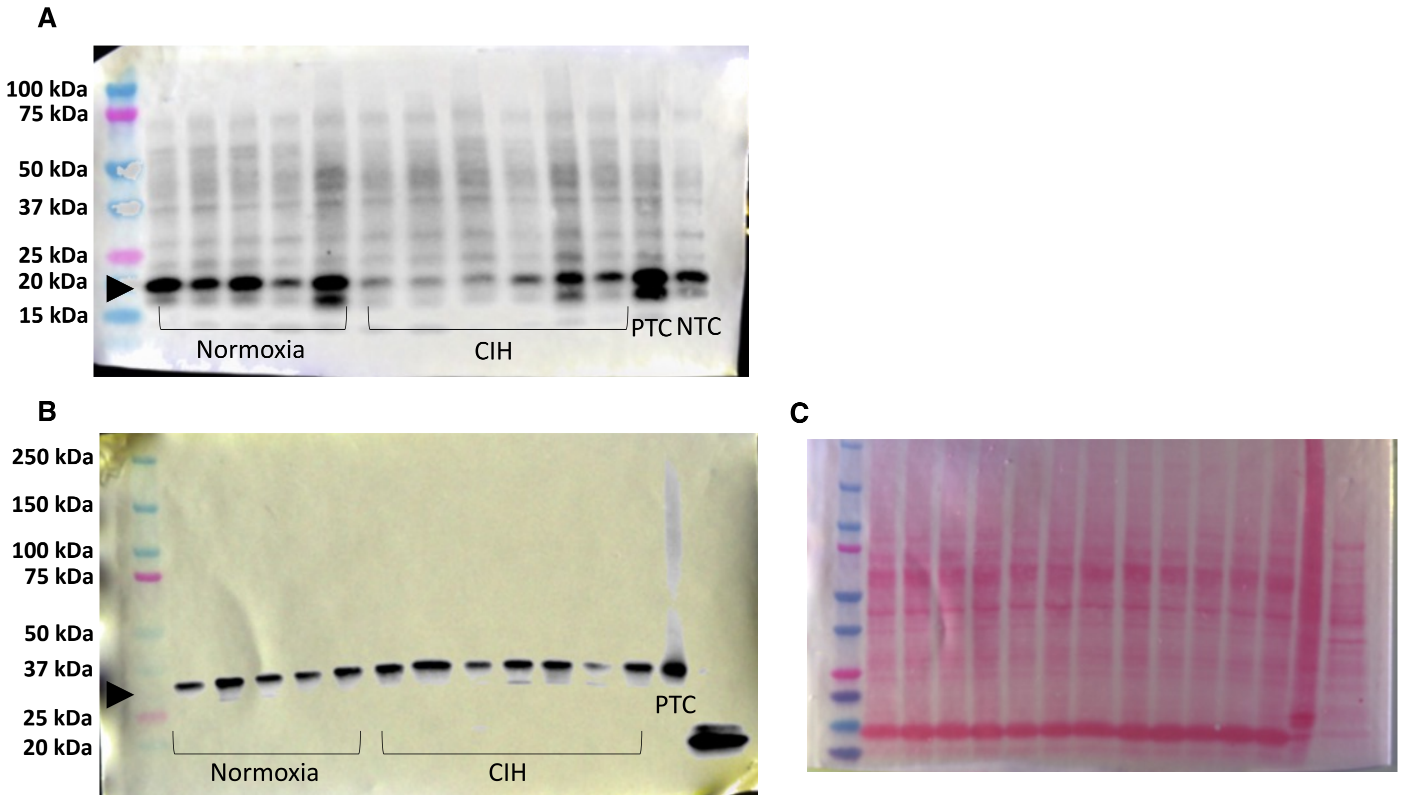


**Figure S5.** Representative immunoblot staining against cleaved-caspase 3 (A), caspase 3 (B) and Ponceau S total protein staining (C) in rat normoxia and chronic intermittent hypoxia (CIH) placentas. Extract of HeLa cells treated with chloroquine was used as positive control (PTC) and HeLa cells untreated was used as negative control (NTC).

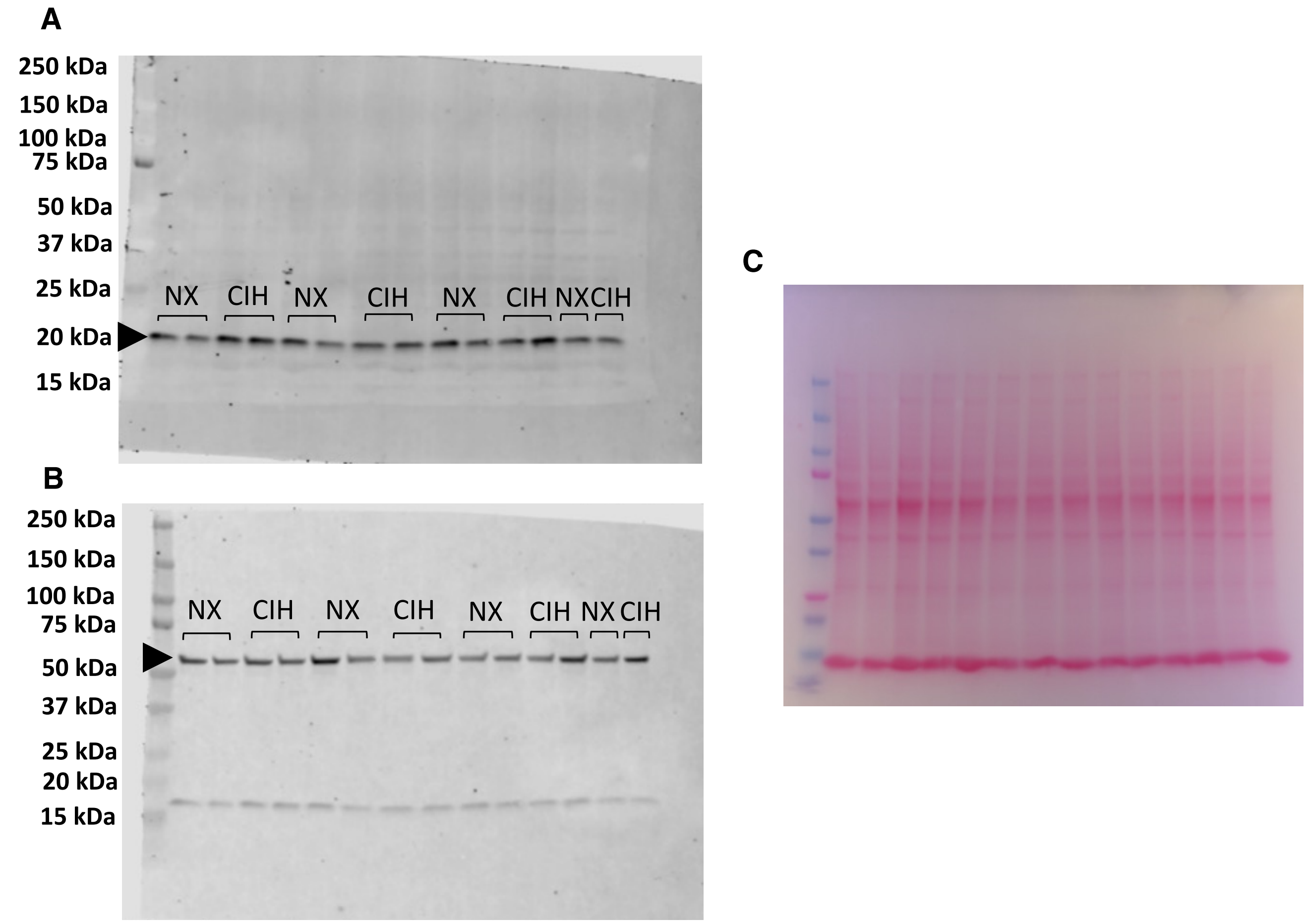


**Figure S6.** Representative immunoblot staining against superoxide dismutase 1 (SOD-1) (A), catalase (B) and Ponceau S total protein staining (C) in rat normoxia (NX) and chronic intermittent hypoxia (CIH) placentas. Antibodies against SOD-1 and Catalase were incubated on the same membrane.

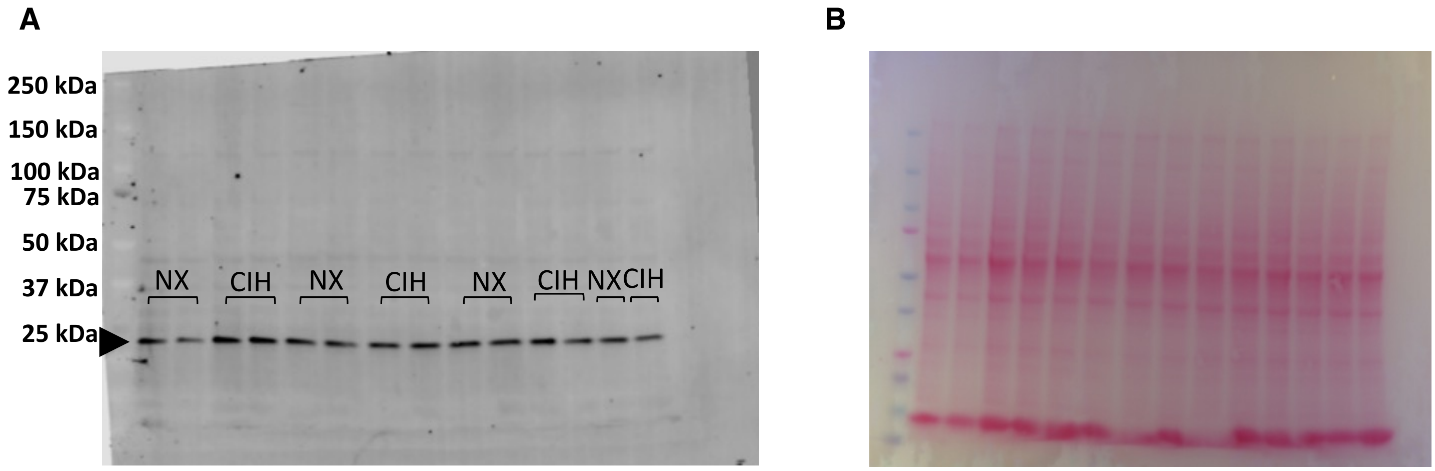


**Figure S7.** Representative immunoblot staining against superoxide dismutase 2 (SOD-2) (A) and Ponceau S total protein staining (B) in rat normoxia (NX) and chronic intermittent hypoxia (CIH) placentas.

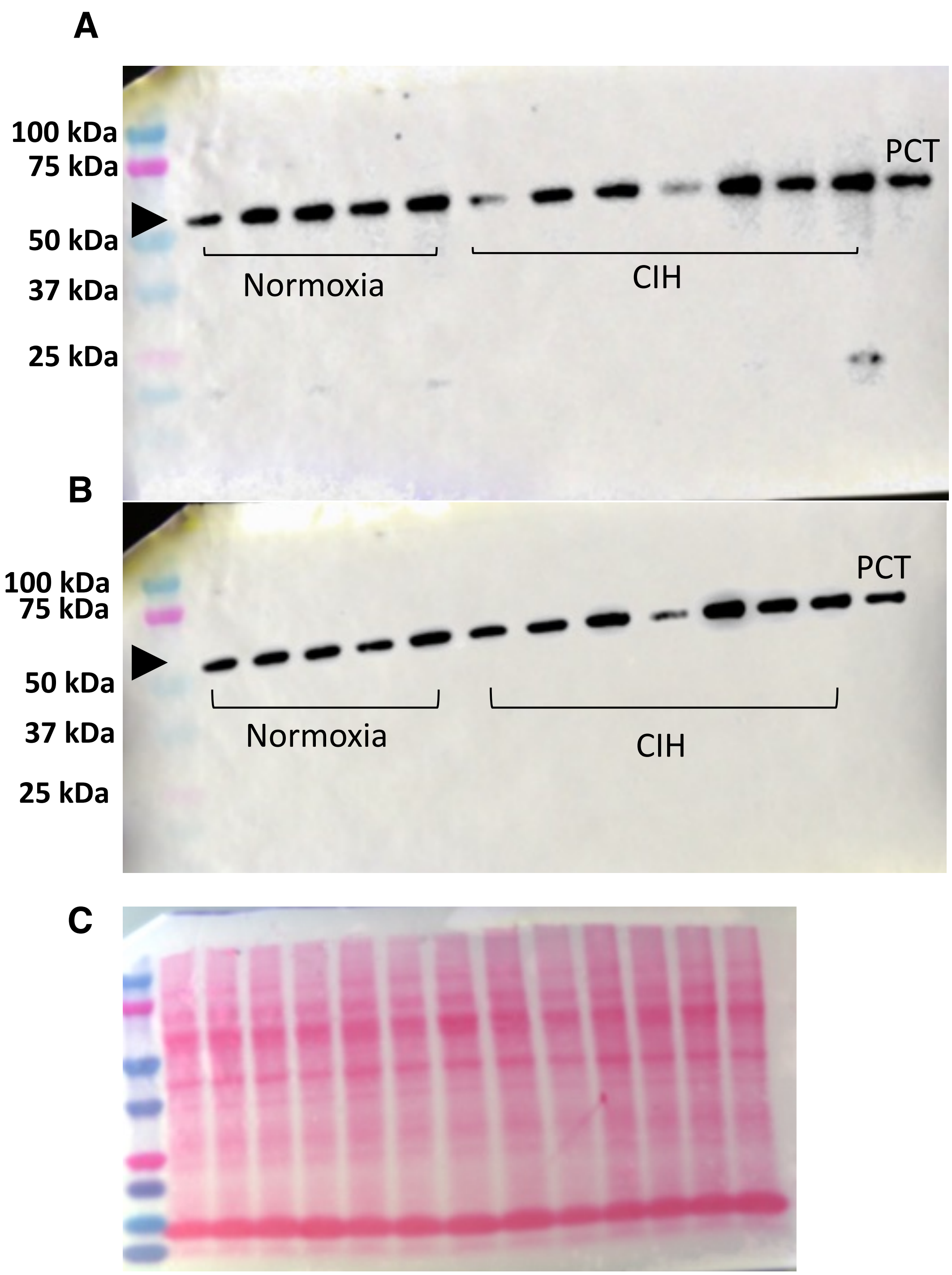


**Figure S8.** Representative immunoblot staining against phospho-Beclin-1 (A), Beclin-1 (B) and Ponceau total protein staining (C) in rat normoxia and chronic intermittent hypoxia (CIH) placentas. Extract of HeLa cells treated with chloroquine was used as positive control (PTC).

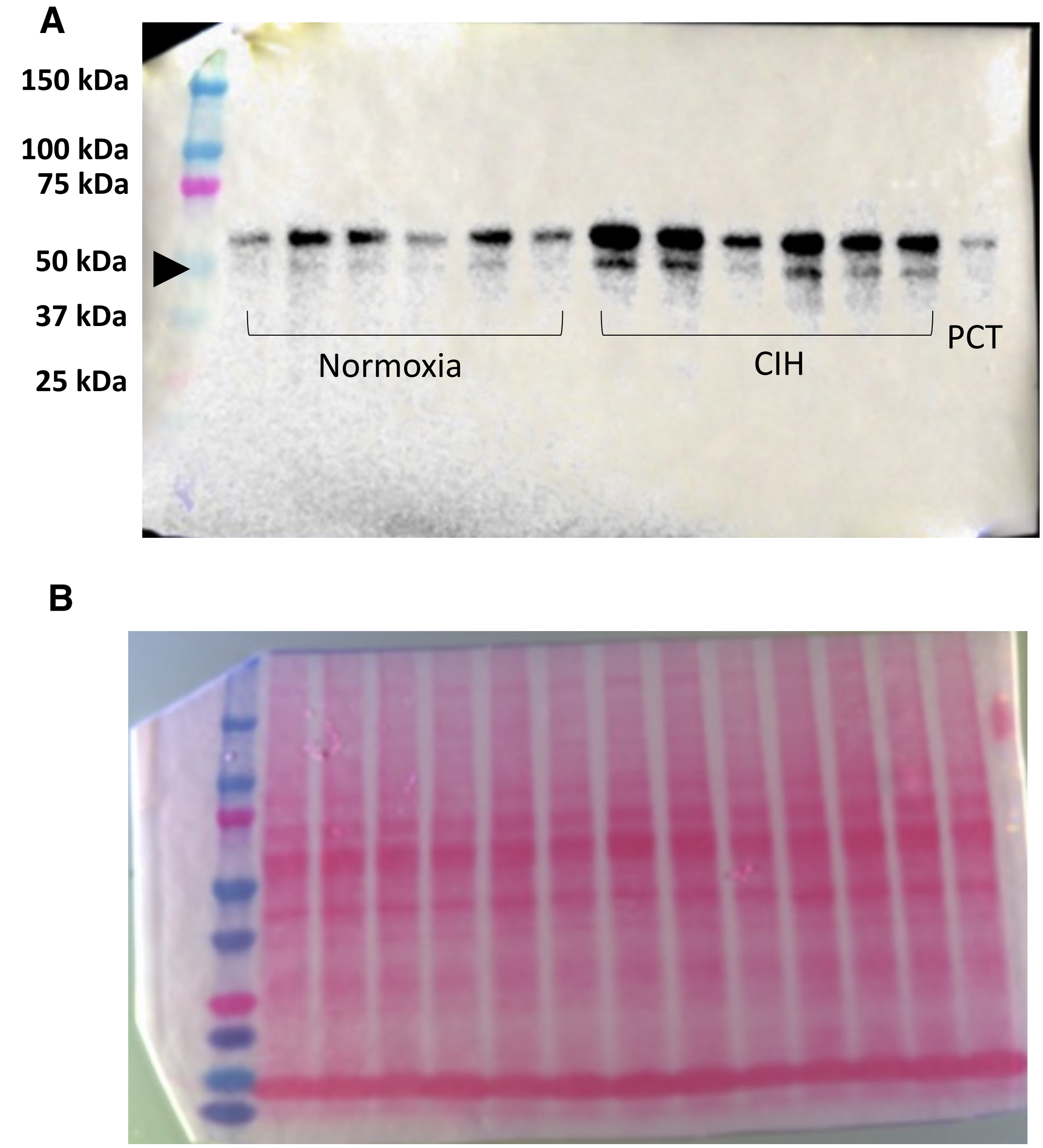


**Figure S9.** Representative immunoblot staining against p62 (A) and Ponceau S total protein staining (B) in rat normoxia and chronic intermittent hypoxia (CIH) placentas. Extract of HeLa cells treated with chloroquine was used as positive control (PTC).

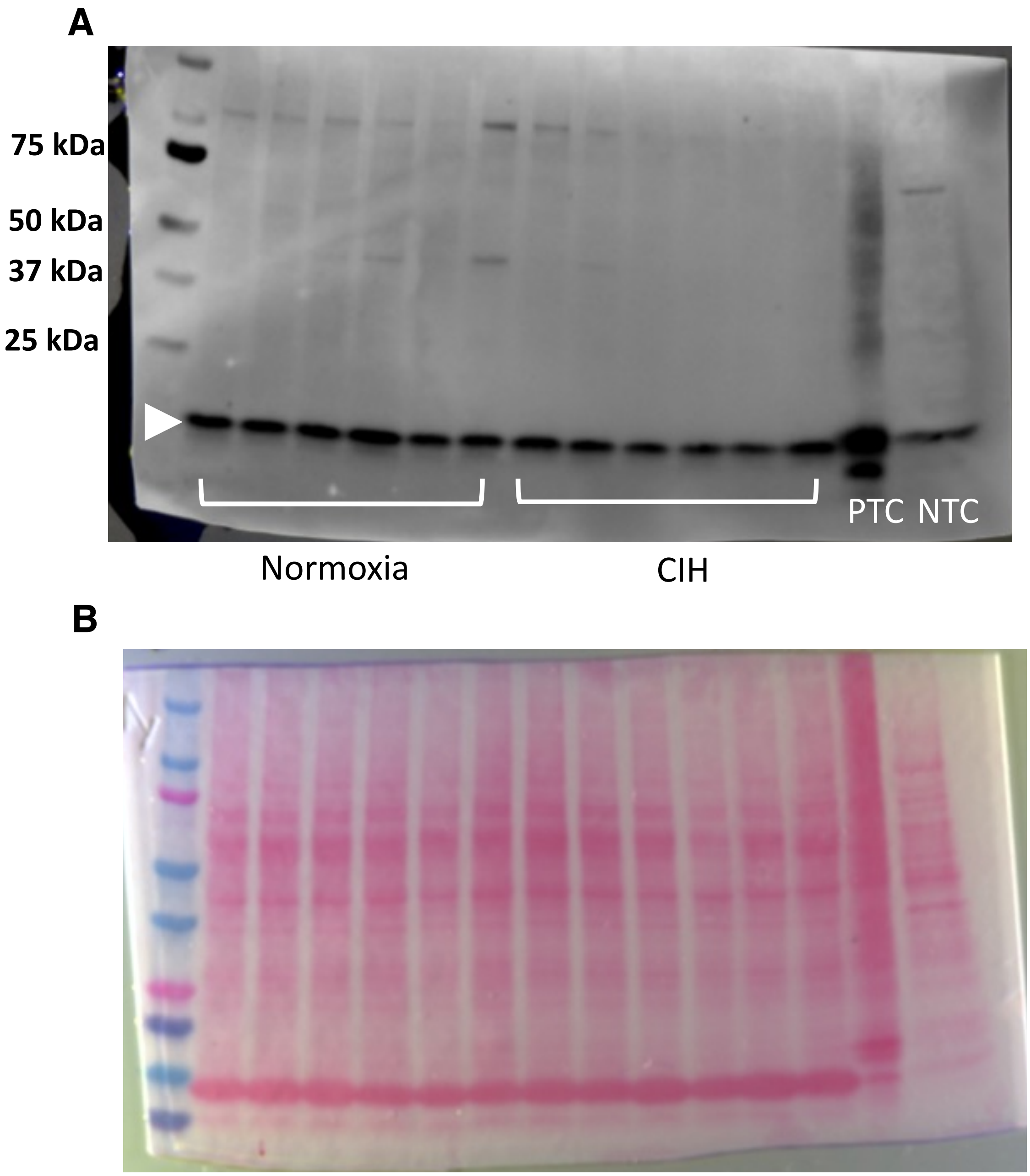


**Figure S10.** Representative immunoblot staining against LC3A/B (A) and Ponceau S total protein staining (B) in rat normoxia and chronic intermittent hypoxia (CIH) placentas. Extract of HeLa cells treated with chloroquine was used as positive control (PTC) and untreated HeLa cells were used as negative control (NTC).

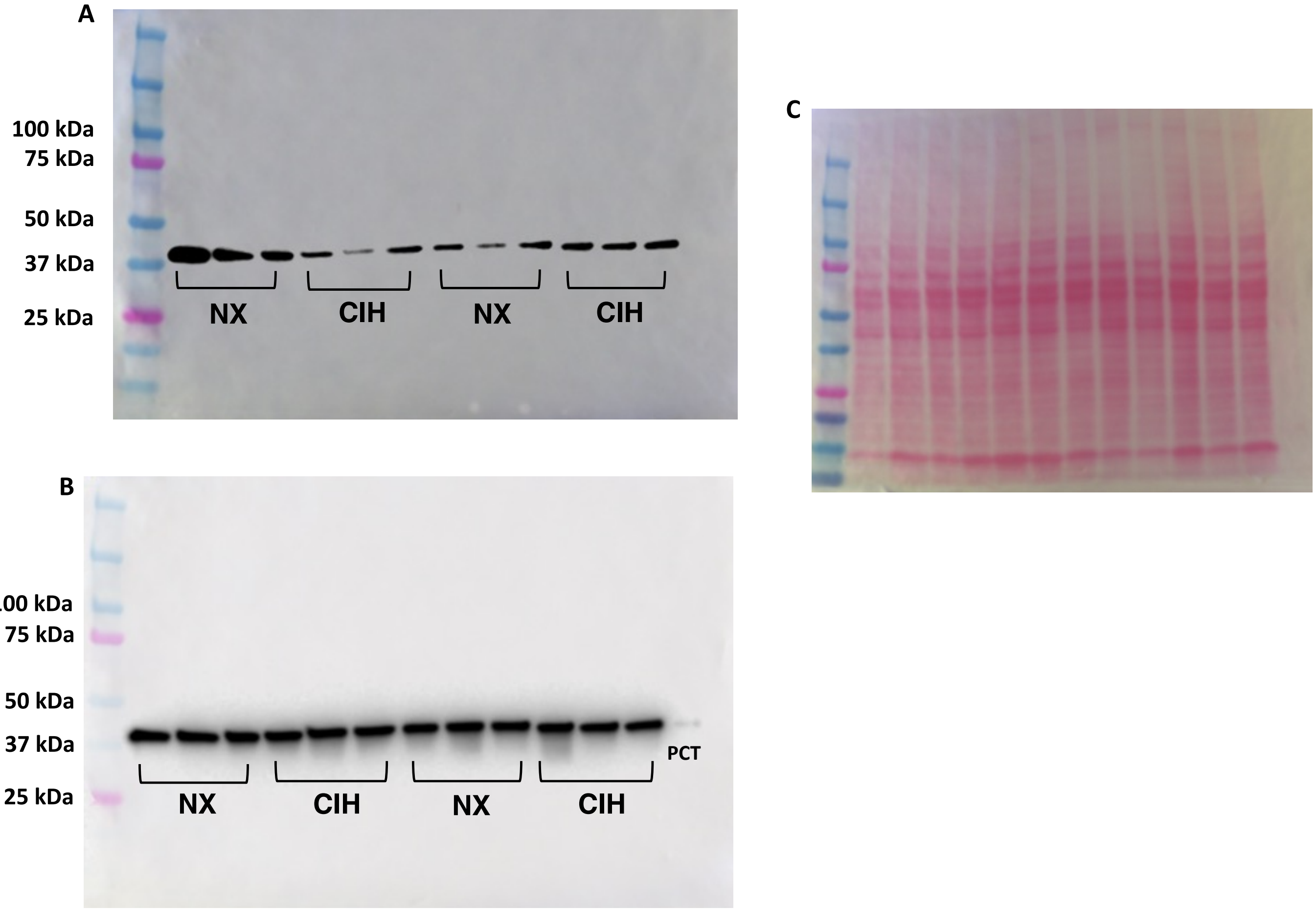


**Figure S11.** Representative immunoblot staining against phospho-p38 (A), p38 (B) and Ponceau total protein staining (C) in rat normoxia (NX) and chronic intermittent hypoxia (CIH) deciduae. C-6 glioma cells treated with anisomycin was used as a positive control (PTC).

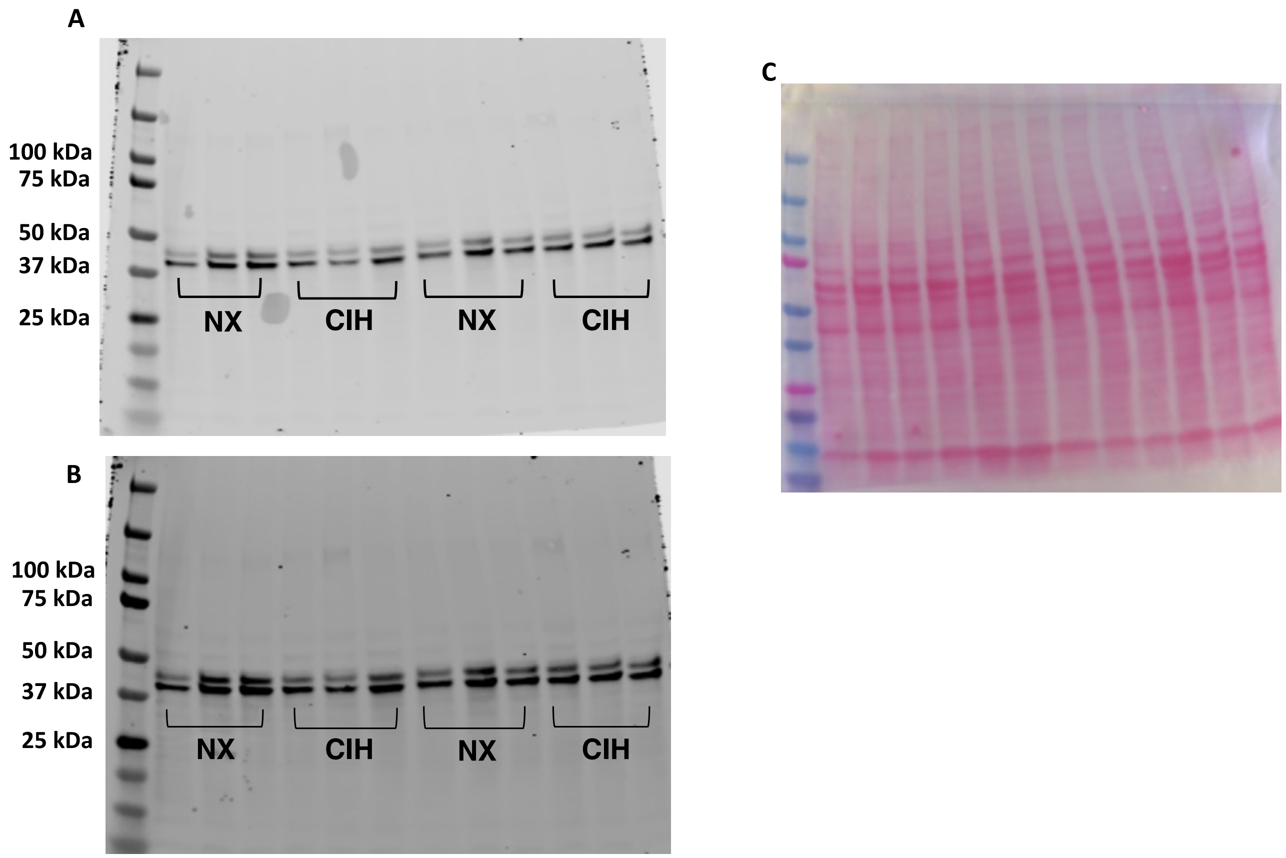


**Figure S12.** Representative immunoblot staining against phospho-ERK 1/2 (A), ERK 1/2 (B) and Ponceau total protein staining (C) in rat normoxia (NX) and chronic intermittent hypoxia (CIH) deciduae.

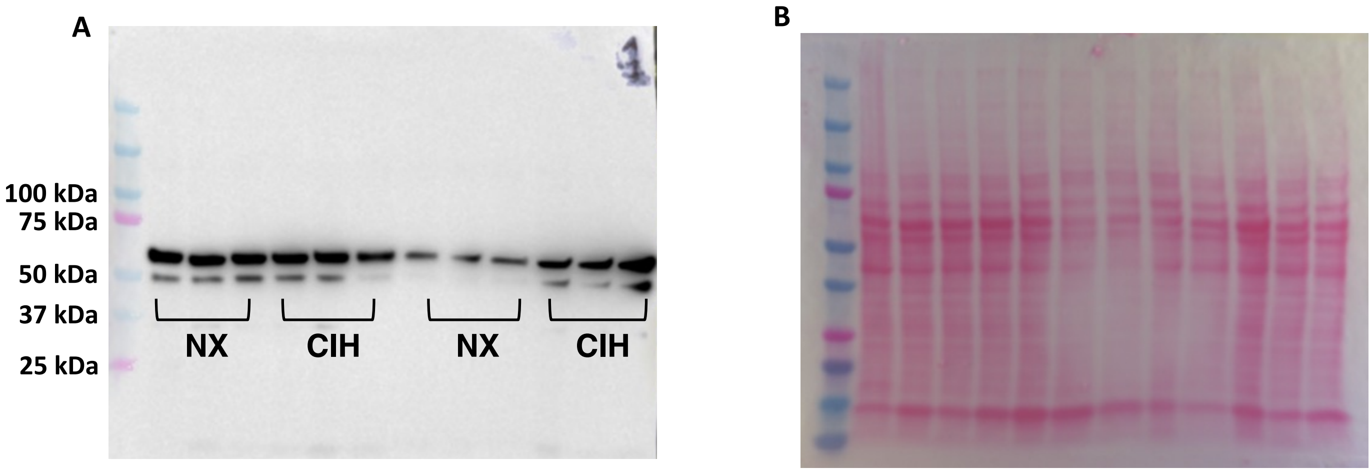


**Figure S13.** Representative immunoblot staining against p62 (A) and Ponceau total protein staining (B) in rat normoxia (NX) and chronic intermittent hypoxia (CIH) deciduae.

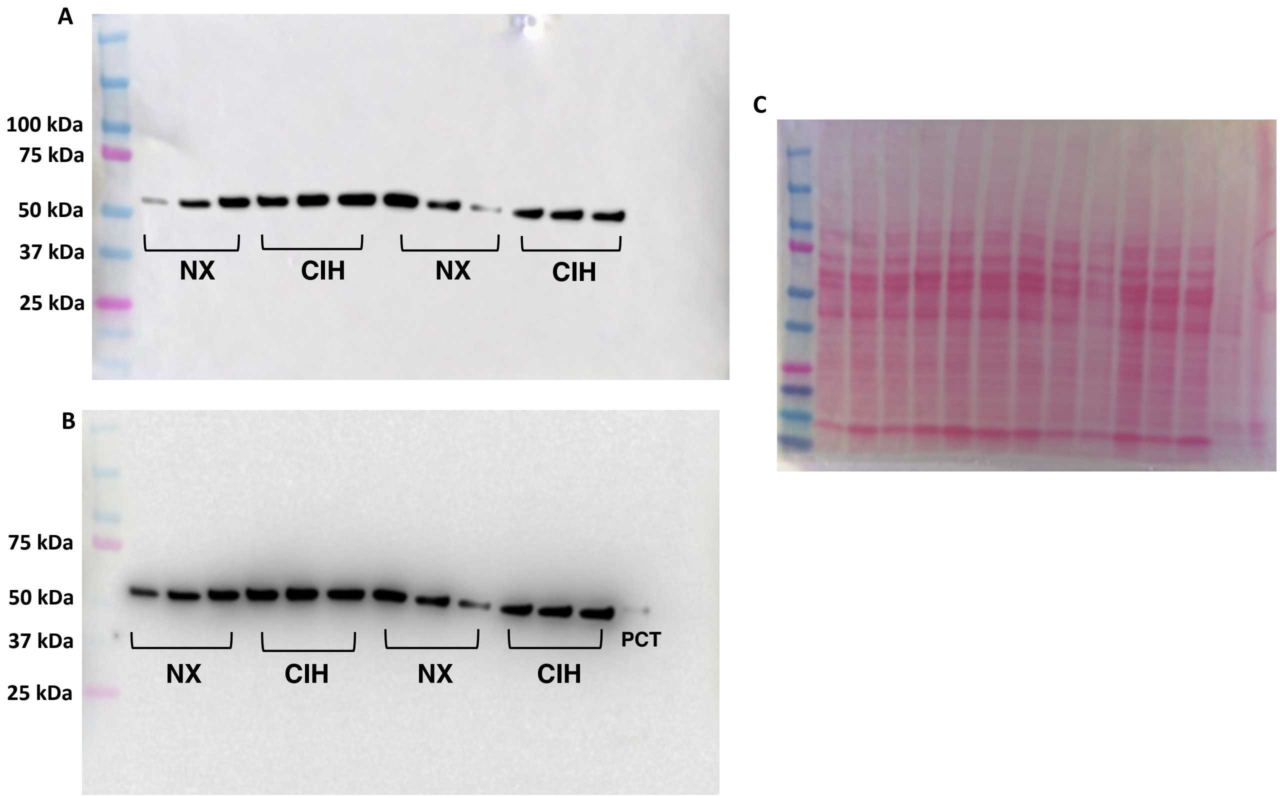


**Figure S14.** Representative immunoblot staining against phospho-Beclin-1 (A), Beclin-1 (B) and Ponceau total protein staining (C) in rat normoxia (NX) and chronic intermittent hypoxia (CIH) deciduae. Extract of HeLa cells treated with chloroquine was used as positive control (PTC).

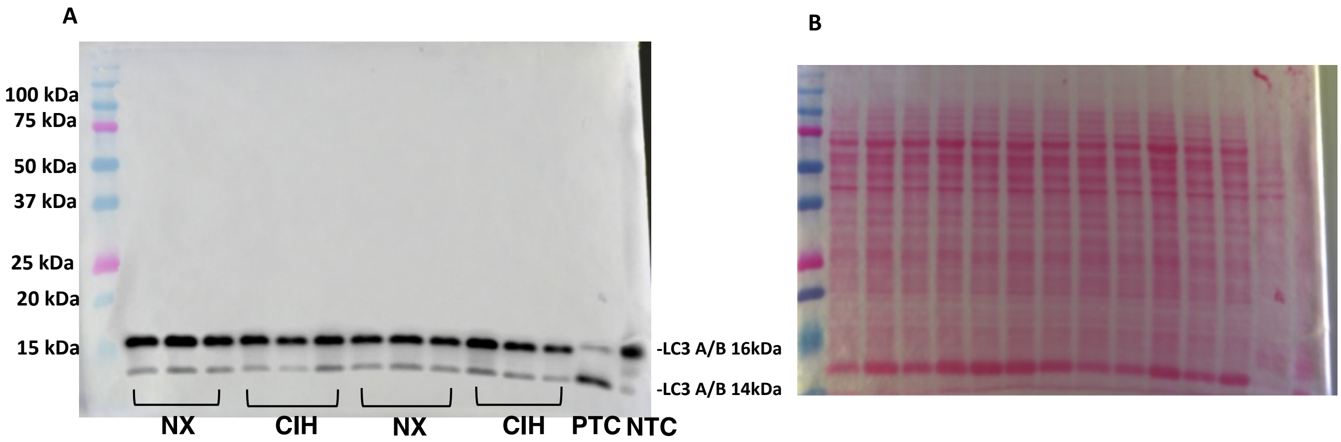


**Figure S15.** Representative immunoblot staining against LC3 A/B(A), and Ponceau S total protein staining (B) in rat normoxia (NX) and chronic intermittent hypoxia (CIH) deciduae. Extract of HeLa cells treated with chloroquine was used as positive control (PTC) and untreated HeLa cells were used as negative control (NTC).

### References

1. **Dhakal P, and Soares MJ**. Single-step PCR-based genetic sex determination of rat tissues and cells. *Biotechniques* 62: 232-233, 2017.
